# Dual Roles of the Conditional Exosomes Derived from *Pseudomonas aeruginosa* Biofilms: Promoting and Inhibiting Bacterial Biofilm Growth

**DOI:** 10.1101/2023.09.06.556615

**Authors:** Marwa Gamal Saad, Haluk Beyenal, Wen-Ji Dong

## Abstract

Antibiotic-resistant biofilm infections have emerged as public health concerns because of their enhanced tolerance of high-dose antibiotic treatments. The biofilm life cycle involves multiple developmental stages, which are tightly regulated by active cell-cell communication via specific extracellular signal messengers such as exosomes. This study was aimed at exploring the roles of *Pseudomonas aeruginosa* exosomes secreted at different developmental stages in controlling biofilm growth. Our results show that exosomes secreted by *P. aeruginosa* biofilms during their exponential growth phase (G-Exo) enhance biofilm growth. In contrast, exosomes secreted by *P. aeruginosa* biofilms during their death/survival phase (D-Exo) can effectively inhibit/eliminate *P. aeruginosa* PAO1 biofilms up to 4.8-log_10_ CFU/cm^2^. The inhibition effectiveness of D-Exo against *P. aeruginosa* biofilms grown for 96 hours improved further in the presence of 10-50 μM Fe^3+^ ions. Proteomic analysis suggests the inhibition involves an iron-dependent ferroptosis mechanism. This study is the first to report the functional role of bacterial exosomes in bacterial growth, which depends on the developmental stage of the parent bacteria. The finding of D-Exo-activated ferroptosis-based bacterial death may have significant implications for preventing antibiotic resistance in biofilms.

**Significance statement:** Antibiotic-resistant bacterial infections caused 1.27 million deaths in 2019 [1], and this number is projected to increase to 10 million deaths annually worldwide by 2050 [2]. Of these infections, up to 80% are caused by biofilm-associated infections [3, 4], which pose a significant challenge to human health. The treatment of biofilm infections remains a formidable problem because of the limited effectiveness of the currently available antibiotics against drug-resistant biofilms [5, 6]. The development of new therapeutic approaches that can effectively combat biofilm infections is required. This study represents a promising solution to antibiotic-resistant biofilm infections. The successful use of exosomes against biofilms opens new possibilities for combating challenging antibiotic-resistant biofilm infections.

## Introduction

Drug-resistant bacterial infections are a significant and growing challenge to global health. Several gram-positive and gram-negative bacterial strains, including *Pseudomonas aeruginosa* and methicillin-resistant *Staphylococcus aureus* (MRSA), have been identified as drug-resistant pathogens, or “superbugs.” In 2019, drug-resistant bacterial infections caused 1.27 million deaths [1], and this number is projected to increase to 10 million annual deaths worldwide by 2050 [2]. The existing antibiotics are often not effective enough at eradicating pathogenic bacteria, making the treatment of such infections a significant challenge and highlighting the urgent need to develop more effective approaches. Antibiotic resistance becomes especially problematic when the pathogenic cells grow as biofilms [7]. Approximately 80% of drug-resistant bacterial infections in the human body are due to biofilms, and bacteria within these biofilms can increase their antibiotic resistance up to 1000-fold [8].

Biofilm development involves multiple stages, including attachment, adaptation followed by exponential growth, maturation, stationary growth and dispersion leading to new biofilm formation. These stages can be influenced by the culture environment, with some stress or hostile cultural conditions forcing the bacterial biofilm into a cell death/survival phase, in which bacterial populations can constantly switch among growth, survival, and death [9]. Each of these developmental stages is tightly regulated by active cell-cell communication via extracellular vesicles (EVs) or exosomes to coordinate cellular processes [10-12].

Exosomes are small membrane-derived EVs with diameters of 30–150 nm that can be secreted by all types of cells. They transfer lipids, proteins, mRNAs, and microRNAs from parental cells to other cells, thus altering the target cells’ behavior, making exosomes important mediators of intercellular communication [13-15]. Exosome-based intercellular communication has been extensively studied in many biological and pathological processes, particularly in the context of cancer [16-18], but it is now recognized as a primordial feature of all living cells [19, 20]. In recent years, it has been recognized that the secretion of exosomes appears to be a conserved process in both gram-negative and gram-positive bacteria. Bacterial exosomes not only engage in various physiological and pathological processes associated with bacteria-host infections but are also involved in regulating essential cellular processes of bacterial life cycles [10, 11], including cellular division and the formation and maintenance of biofilms [12]. We hypothesized that the role of bacterial exosomes in biofilm development is culture condition dependent, i.e., that the exosomes secreted by biofilms in their exponential growth phase and in their survival/death phase would promote biofilm growth and inhibit biofilm growth/formation, respectively.

To test our hypothesis, we investigated whether exosomes derived from *P. aeruginosa* (PAO1) biofilms at different developmental stages can be utilized to control planktonic growth and biofilm formation. These stage-dependent exosomes are referred to as conditional bacterial exosomes. Briefly, we extracted exosomes released by PAO1 biofilms at their growth stage (G-Exo) and their survival/death phase (D-Exo), respectively. We then examined how these conditional exosomes control PAO1 biofilm growth under different conditions. The results of our study demonstrate the potential of the conditional bacterial exosomes as alternative antibiotic agents for addressing the issue of antibiotic resistance currently facing society.

## Results

### 1. *Isolation and characterization of conditional exosomes secreted by* P. aeruginosa *PAO1 biofilms*

**Fig. 1** shows *P. aeruginosa* PAO1 biofilm growth curves at 33°C and 37°C. When growing at 37°C, *P. aeruginosa* PAO1 biofilm reached a steady cell number of 7.7 CFU/cm^2^ at the 14th hour and the plateau phase lasted to the 129^th^ hour. However, the death/survival phase was not distinguishable before the 120th hour. Under a 33°C culture condition, PAO1 biofilm growth clearly showed four growth stages: lag, exponential growth, plateau, and survival/death, with a steady cell number of 6 CFU/cm^2^ in the plateau phase. Based on these observations, in this study, all conditional exosomes were extracted from *P. aeruginosa* PAO1 biofilms cultured under the 33°C condition. However, all functional tests of the obtained exosomes on *P. aeruginosa* PAO1 biofilm growth were performed under the 37°C culture condition.

**Fig. 1.**
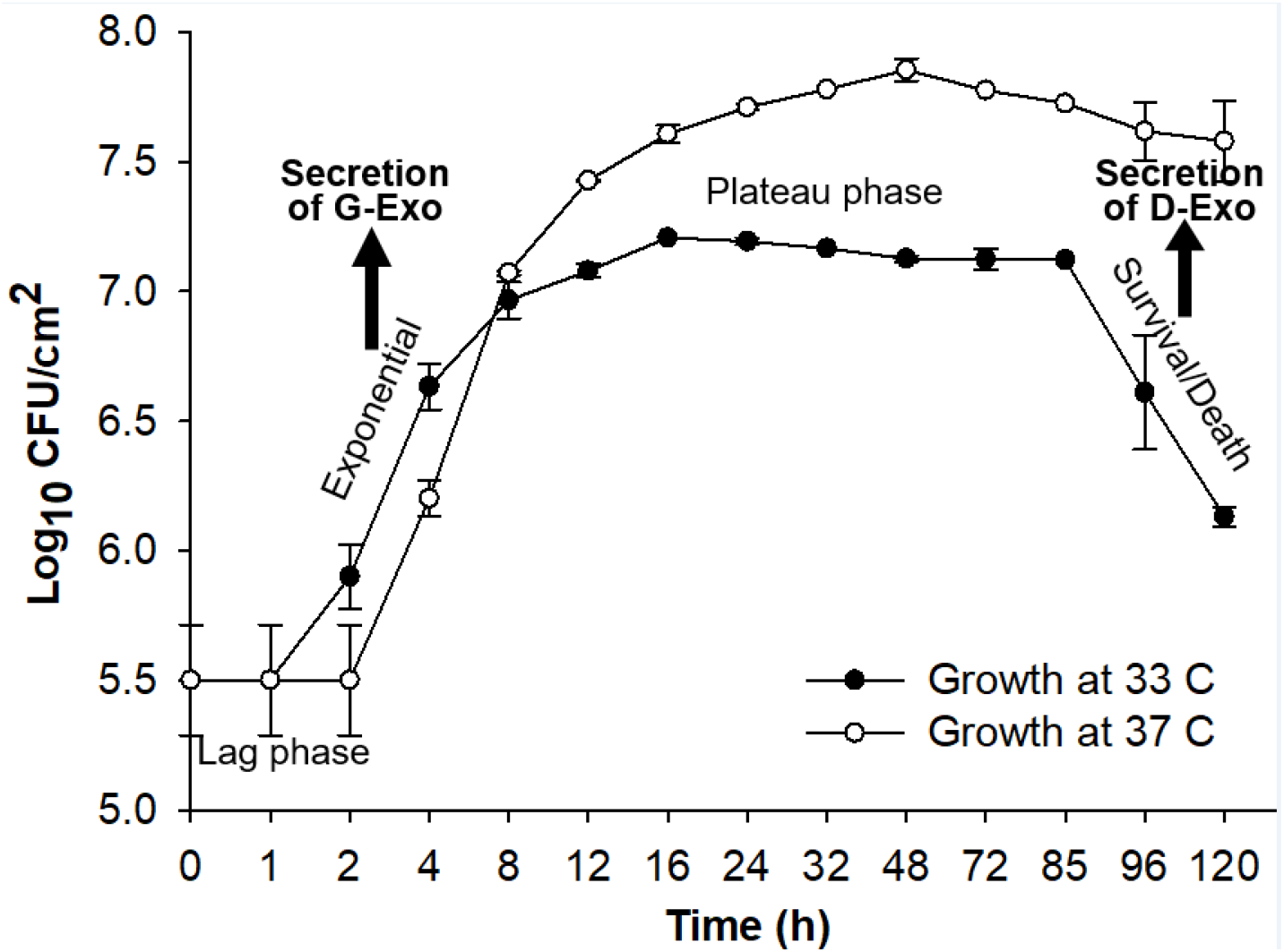
Growth curves of *P. aeruginosa* PAO1 biofilms inoculated incubated at 32.7°C and 37°C, respectively on TSA plates. Four stages of biofilm growth are clearly presented for the growth curve of 32.7 °C. Also indicated are the stages where the biofilms were collected to extract G-Exo and D-Exo, respectively. Note: the scale of the x-axis is not linear. Data points indicate mean biomass, and error bars indicate standard deviations (n = 4).

G-Exo or D-Exo of *P. aeruginosa* PAO1 biofilms was extracted from biomasses of the biofilms grown at 33°C. The biomasses were collected at specific developmental stages, as indicated in **Fig. 1**, using conventional differential centrifugation protocols [21-24], which are given in the Materials and Methods section and *<u>Supplementary Information section S1</u>* (**Fig. S1**). Using this protocol, a total of 874.1 ± 20 μg exosomal proteins could be extracted from a biofilm-covered membrane paper with a cell concentration of 4.4x10^5^ ± 7.3 x10^2^ CFU/cm^2^.

To correlate the protein concentration of exosomes, stained exosome samples of different protein concentrations were analyzed using confocal laser scanning microscopy (CLSM) (see *<u>Supplementary Information section S2</u>*). The results (shown in **Fig. S2** and **Fig. S3**) established the correlation between the protein concentrations of the exosomes and their average particle numbers in a solution, with every exosome particle equivalent to an average exosomal protein content of 0.00188± 8x10^-4^ μg/ml.

The shapes and sizes of the purified G-Exo and D-Exo were characterized using transmission electron microscopy (TEM) analysis (see *<u>Supplementary Information section S3</u>* and **Fig. S4**). The size of exosomes extracted during the exponential growth phase was larger (112.9 ± 3.7 nm) than that of exosomes extracted during the death/survival phase (33.2 ± 0.9 nm). These results demonstrate that *P. aeruginosa* PAO1 biofilms release different populations of exosomes at different stages, consistent with results observed for tumor cells and HeLa cells, which release different populations and subpopulations of EVs [25, 26]. This suggests that these distinct populations of exosomes secreted by *P. aeruginosa* PAO1 biofilms at different stages have different functional roles in the biofilm life cycle.

### 2. The functional effects of G-Exo and D-Exo on the growth of P. aeruginosa PAO1 biofilm

To investigate our hypothesis that exosomes produced by a bacterial biofilm at a specific growth stage can influence the growth of recipient bacterial communities in the same developmental direction, we conducted a disc diffusion assay (described in the Materials and Methods section) using both D-Exo and G-Exo to examine their functional effects on the growth behavior of *P. aeruginosa* PAO1 biofilm (see *<u>Supplementary Information section S4</u>* and **Fig. S5**). We used PBS buffer and the antibiotic tetracycline as negative and positive controls, respectively, to assess the efficacy of D-Exo/G-Exo in inhibiting/promoting the formation and growth of *P. aeruginosa* PAO1 biofilms in comparison to tetracycline. Given that *P. aeruginosa* PAO1 is a known antibiotic-resistant superbug, we used a sub-inhibitory concentration of tetracycline (1 μg/μL) in the test to ensure a clear zone on the dish (**Fig. S5A**). This is much higher than the medically tolerant doses (0.033-0.22 μg/μL) used for adult admissions [27]. Although the qualitative nature of the diffusion test made it difficult to discern the role of G-Exo in biofilm formation and growth, our D-Exo exhibited effective and dose-dependent inhibition of *P. aeruginosa* PAO1 biofilm growth across a range of exosomal protein concentrations (0.11, 0.22, and 0.33 μg/μL).

In order to confirm the importance of intact exosome structure for the observed effects, we conducted an experiment in which we disrupted the membranes of D-Exo and tested whether this affected their ability to inhibit biofilm growth. We achieved this by subjecting the D-Exo to water bath sonication, which lysed the exosomes (see *<u>Supplementary Information section S6A</u>*). We then used the sonicated D-Exo in the diffusion susceptibility test and compared the results to those obtained with intact D-Exo (see *<u>Supplementary Information section S6B</u>*). We found that the sonicated D-Exo had a much fainter inhibitory effect on biofilm growth than intact D-Exo (**Fig. S5B)**, indicating that an intact exosomal structure is essential for exosomes to inhibit biofilm growth. These qualitative results are the first to demonstrate the potential of exosomes secreted by a bacterial biofilm in its death/survival phase for use against bacterial biofilm growth of the same bacterium.

To assess how D-Exo and G-Exo affect biofilm growth, we monitored *P. aeruginosa* PAO1 biofilm growth in terms of Log_10_CFU/cm^2^. The functional efficacy of the exosomes varied depending on the maturity of the biofilm at each stage of development. **Fig. 2A** illustrates the growth of *P. aeruginosa* PAO1 bacterial biofilms grown for 2 h and treated with D-Exo and G-Exo compared to biofilms treated with PBS as a control. The results show that, compared to the control, the presence of G-Exo at a protein concentration of 0.028 μg/μL significantly increased the growth rate of the biofilms, while D-Exo at a protein concentration of 0.33 μg/μL showed a bacteriostatic effect on biofilm formation and growth. **Fig. 2B** shows the effects of D-Exo/G-Exo on the growth of *P. aeruginosa* PAO1 biofilms formed after 8 hours of growth (the exponential growing phase) after bacterial inoculation. It reveals a 2-log_10_ increase in the biomass of the biofilm after one dose of G-Exo at a protein content of 0.037 μg/μL. In contrast, treatment of the biofilm with one dose of D-Exo at a protein content of 0.33 μg/μL reduced the biofilm biomass by 4.8-log_10_ compared to the control. These experiments demonstrated that one-dose treatment of D-Exo/G-Exo significantly affected the behavior of biofilms formed in the initial lag and exponential growth phases (2-h and 8-h biofilms). The results from the D-Exo experiments suggest that D-Exo can be used to treat *P. aeruginosa* PAO1 biofilms at the early stage of biofilm growth. These quantitative results provide direct evidence supporting our hypothesis that G-Exo promotes biofilm growth and D-Exo inhibits biofilm formation and growth.

**Fig. 2.**
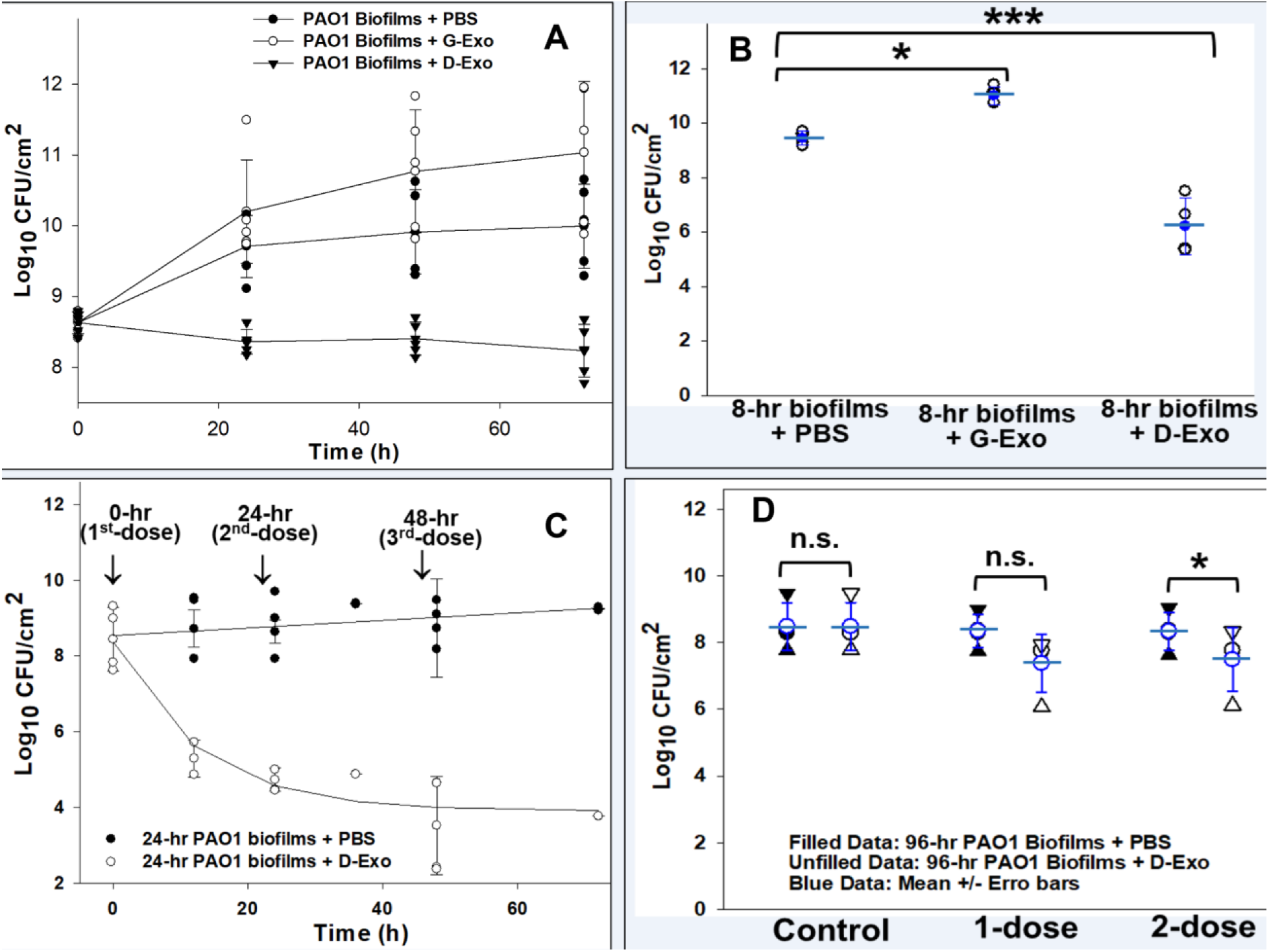
Functional effect of G-Exo and D-Exo on the growth behavior of *P. aeruginosa* PAO1 biofilms. **A**: Growth of *P. aeruginosa* PAO1 biofilms starting from the initial phase after one-dose treatment with G-Exo (0.028 μg/μL) and D-Exo (0.33 μg/μL), respectively. The data points are the times at which the biomass of the biofilms under each condition was collected and analyzed. **B**: Growth of *P. aeruginosa* PAO1 biofilms (8-h) formed during the log growth phase after one-dose treatment with G-Exo (0.037 μg/μL)orD-Exo (0.33 μg/μL). Biomasses were collected and analyzed 24 hours after the biofilms were treated. **C**: Growth of *P. aeruginosa* PAO1 biofilms (24-h) developed during the stationary plateau phase after multidose treatment with D-Exo (0.33 μg/μL). Three doses ofD-Exo were applied to the biofilms— at 0, 24 and 48 h (indicated by **↓** signs). The data points on each curve are the times at which the biomass of the biofilms under each condition was collected and analyzed. **D**: Growth of *P. aeruginosa* PAO1 biofilms (96-h) developed during the survival/death phase after one dose or 2 doses with a 12-h interval of D-Exo (0.33 μg/μL) or PBS. The mean value and error bar of each group of data are given in blue. For all data points, the biomasses of biofilms were collected and analyzed 24 hours after each treatment. P value <0.05 (*) and <0.01 (***).

Bacterial biofilms are notoriously difficult to treat with antibiotics, as they become increasingly resistant as they mature, particularly in the stationary plateau phase. In this study, we aimed to determine whether D-Exo could effectively treat mature biofilms in the stationary phase. **Fig. 2C** shows the inhibitory effects of multiple sequential doses of D-Exo at a protein concentration of 0.33 μg/μL on 24-hour biofilms in the plateau stage. Biomass analysis of the treated biofilms 24 hours after the first dose showed a decrease in biomass of more than 3-log_10_ compared to the control. After further doses at 24 hours and 48 hours, respectively, biofilm growth was inhibited by an additional 1.5-log_10_. The observation of a total inhibition effect of 4.8-log_10_ suggests that a multi-dose approach using D-Exo could be used to treat mature biofilms.

After once reaching the death/survival phase under a slow nutrition depletion condition, biofilms live in a cryptic growth mode and secrete D-Exo to coordinate bacterial functions to maintain a small core of the population alive by gradually recycling nutrients derived from dead cells. In this growth mode, biofilms become aged biofilms that are hard to kill and more resistant to environmental stresses including antibiotics [28, 29]. In this experiment, we further investigated the effects of D-Exo on *P. aeruginosa* PAO1 biofilms (96-h) by applying either one dose of D-Exo at a protein concentration of 0.33 μg/μL directly to the aged biofilms or two doses separated by a 12-h interval. Analysis of the biomass collected 24 hours after the D-Exo treatments is shown in **Fig. 2D**. Since the D-Exo was extracted from the same populations of the biofilms, as expected, treatment with 2 doses of D-Exo only reduced the population of the aged biofilms by <1-log_10_.

### 3. Interactions between the exosomes and bacterial cells

To investigate whether the observed effects of D-Exo/G-Exo on *P. aeruginosa* PAO1 biofilm growth are due to direct interactions between the exosomes and target cells, we used TEM and CLSM to image potential interactions. The TEM images in **Fig. 3A** show that the G-Exo attached to the membrane surface of a target cell, triggering structural changes that led to exosomal uptake. In contrast, **Fig. 3C** shows that incubation with D-Exo led to deterioration of the bacterial cellular wall and subsequent cell death. These results are supported by CLSM experiments. **Fig. 3B** shows that *P. aeruginosa* PAO1 cells became green fluorescent after being incubated with DAPI-labeled G-Exo, indicating uptake of the DAPI-labeled G-Exo by the bacteria. To image bacterial cell viability, the bacterial cells were positively stained with live/dead BacLight Bacterial Viability Kits. **Fig. 3D** shows that most bacterial cells in control samples (no D-Exo presence) were alive and emitting green fluorescence with a total cell fluorescence of 2.5x0^5^ ± 2.9x10^3^; however, after incubation with D-Exo, most of the cells were dead and emitting red fluorescence, indicating cell death; the green living cell fluorescence decreased to 3.1x10^3^ ± 1.4x10^2^. Overall, these findings suggest that the observed distinct features of communication between bacterial cells and G-Exo and D-Exo are directly related to the roles of G-Exo and D-Exo in promoting and inhibiting bacterial growth, respectively.

**Fig. 3.**
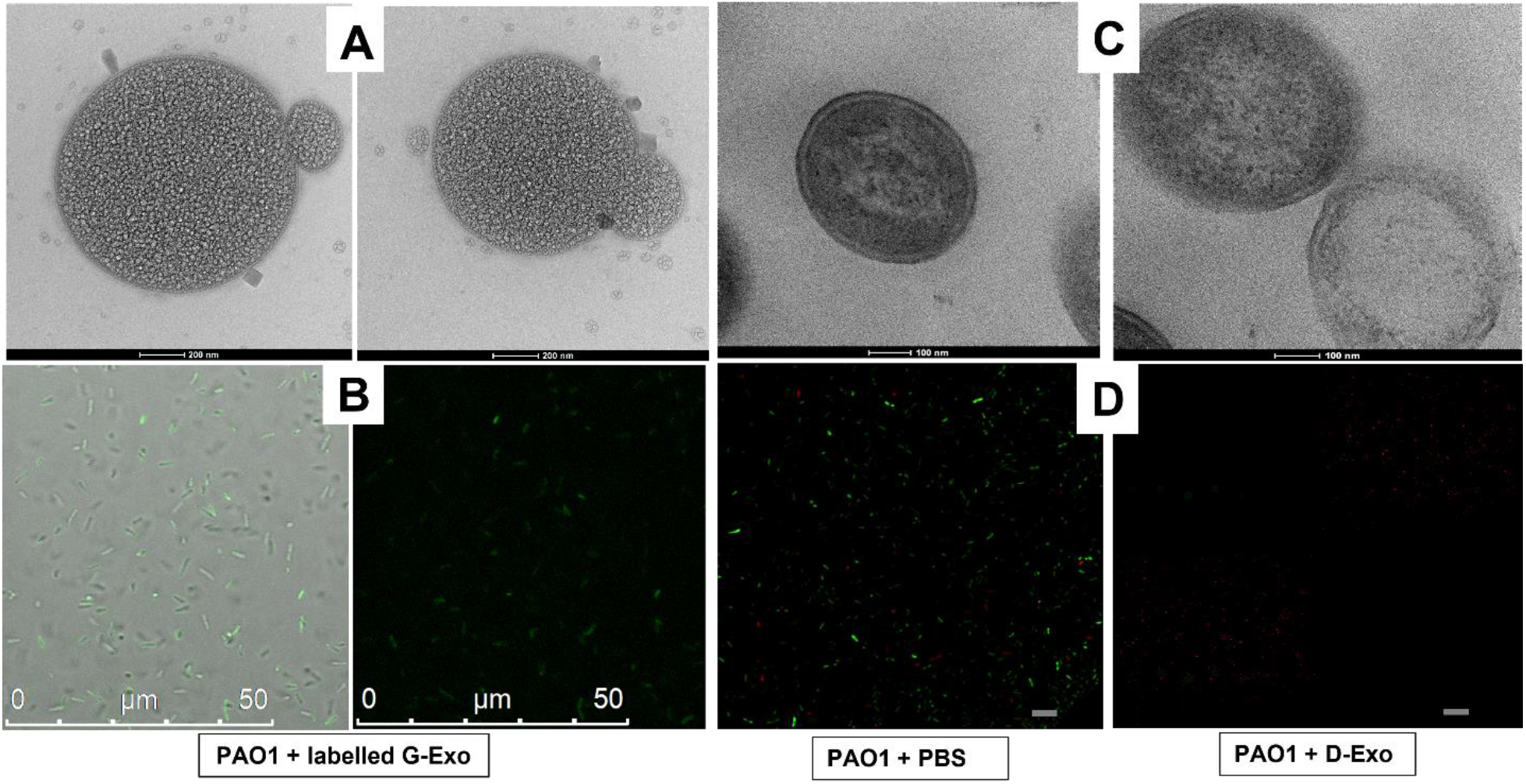
TEM and confocal laser scanning microscopy (CLSM) of the interactions of the G-Exo/D-Exo and *P. aeruginosa* PAO1 bacterial cells. **A**: TEM image of the bacterial cell after incubation with G-Exo added to the cell culture. After 24 h, the cells were negatively stained with 2% uranyl acetate before TEM images were taken (magnification power = 60000X). **B:** CLSM images of the bacterial cells that were incubated with DAPI-dye-labelled G-Exo. The images were taken with an excitation of 359 nm. Under this UV excitation, the recipient cells of DAPI-labeled G-Exo emit green fluorescence [64]. The image on the left shows an overlay of the brightfield and fluorescent images, while the image on the right shows the fluorescent image only. The g reen fluorescence indicates the presence of the DAPI-labeled G-Exo. **C**: TEM image of a bacterial cell in the log phase after incubation with D-Exo. The cells were negatively stained with 2% uranyl acetate before TEM images were taken (magnification power = 40000X). **D:** CLSM images of the bacterial cells that were positively stained with LIVE/DEAD BacLight Bacterial Viability Kits (excitation/ emission: 480/500 nm for SYTO 9 and 490/635 nm for propidium iodide). The <u>image on the left</u> shows the greenly fluorescent live cells treated with PBS buffer; the <u>image on the right</u> shows the red dead cells (scale = 75 μm) after incubation with D-Exo.

### 4. Proteomic analysis of the G-Exo and D-Exo

The proteomic profiles of the G-Exo and D-Exo populations are presented in **Table S1** and **Table S2**, respectively. A total of 93 and 88 highly abundant internal proteins were identified in the G-Exo and D-Exo samples, respectively. The protein profiles of these two populations of bacterial exosomes were found to be significantly different. The proteins carried by G-Exo were primarily associated with cell division and biosynthesis, while those carried by D-Exo were mainly involved in inhibiting cell growth, ROS production and iron acquisition to promote cell survival or induce cell death.

When the G-Exo and D-Exo interacted with the surfaces of the same *P. aeruginosa* PAO1, they had different effects on the cells: the G-Exo was taken up by the cells (**Fig. 3A**) and caused increased cell growth (**Fig. 2A and 2B**), while the D-Exo caused cell rupture (**Fig. 3C**) and cell death (**Fig. 2B and 2C**). To understand how the surface proteins of the G-Exo and D-Exo contributed to their unique roles we identified key proteins that were uniquely present in different abundances on the surfaces of the D-Exo and G-Exo. These proteins are listed in **Table 1**. Among them, eight proteins (**#1-#8**) were exclusively present on the D-Exo surface, while two (**#9** and **#10**) were exclusively present on the G-Exo surface. Protein **#9** on the G-Exo surface is involved in the sulfate transport system of bacteria, which is responsible for energy coupling to the transport system, while protein **#10** belongs to the CusCFBA copper efflux system, which plays a crucial role in copper homeostasis within the bacteria [30]. With these important functional proteins on its surface, the G-Exo may enhance bacterial viability and promote cell growth under normal culture conditions through interactions with target bacterial cells.

**Table 1.** Distribution of key surface proteins on G-Exo and D-Exo from PAO1 biofilms.

|  |  |  |  |  |
| --- | --- | --- | --- | --- |
| No. | Proteins | Possible functions | G-Exo | D-Exo |
| 1 | Heme/hemoglobin uptake outer membrane receptor PhuR | Promote acquisition of heme as iron resource for bacterial growth. Because of its capability of transferring electrons at physiological pH, iron plays a critical role in bacterial physiology as an essential component of metabolic enzymes and regulatory proteins. | x | H |
| 2 | Hemin receptor |  | x | H |
| 3 | Putative outer membrane ferric siderophore receptor |  | x | H |
| 4 | Putative hydroxamate-type ferrisiderophore receptor |  | x | H |
| 5 | Heme acquisition protein HasAp |  | x | H |
| 6 | TonB-dependent receptor | Either mediates the release of iron into the periplasm of the bacteria or transports ferric enterobactin into the periplasm. | x | H |
| 7 | Putative TonB-dependent receptor |  | x | H |
| 8 | Ferric enterobactin receptor |  | x | H |
| 9 | Sulfate/thiosulfate import ATP-binding protein CysA | The sulfur-regulated gene (CysA) that encodes *the membrane-associated ATP-binding protein of the sulfate transport system of bacteria is responsible for energy coupling to the transport system.* | H | x |
| 10 | HlyD_D23 domain-containing protein | The membrane fusion proteins of the CusCFBA copper efflux system that play a crucial role in copper homeostasis within the bacteria, which is directly associated with bacterial viability. | H | x |
| **Note:** H – high abundancy; x – not found. |  |  |  |  |

The eight D-Exo surface proteins can be categorized into two groups. Proteins **#6-#8** are membrane receptors that transport ferric enterobactin into the periplasm or release iron into the periplasm of the bacteria. The second group of D-Exo surface proteins (**#1-#5**) is membrane receptors or functional proteins that promote bacterial growth by enhancing the acquisition of iron resources. Iron is an essential component of metabolic enzymes and regulatory proteins that support the growth and survival of most bacterial species. However, excessive uptake of iron by bacteria can be detrimental because of the iron-triggered Fenton/Haber-Weiss reaction, which produces harmful ROS such as superoxide (O^2−^), hydrogen peroxide (H_2_O_2_), and the highly destructive hydroxyl radical (^·^OH). Accumulation of ROS can inactivate key functional enzymes and activate bacterial programmed cell death [31-38].

### 5. Mechanistic investigations of inhibition effects of the D-Exo on biofilm growth

Based on these proteomic data, we hypothesized that D-Exo induces excessive iron uptake by the D-Exo recipient cells, resulting in the accumulation of ROS and ultimately the activation of bacterial cell death. This hypothesis was tested by examining how the poor efficacy of the D-Exo against biofilms grown for 96 h (**Fig. 2D**) was improved in the presence of ferric ions. Compared to the PBS control, the presence of ferric ions alone marginally promoted biofilm growth, from 8.2-log_10_ to 8.7-log_10_ (**Fig. S6B**), while 2-dose treatments of D-Exo alone only reduced the aged biofilm growth by less than 1-log_10_ (**Fig. S6A**). However, when ferric ions were applied to the 96-h *P. aeruginosa* PAO1 biofilms together with D-Exo, the growth of the biofilms was significantly inhibited (**Fig. 4**), and the inhibition efficiency was found to be Fe^3+^ concentration dependent. As [Fe^3+^] increased from 10 μM to 50 μM, the inhibition of biofilm growth increased from a 2.6-log_10_ reduction to a 3.4-log_10_ reduction. These results demonstrate that both D-Exo and ferric ions are needed to treat the biofilms.

**Fig. 4.**
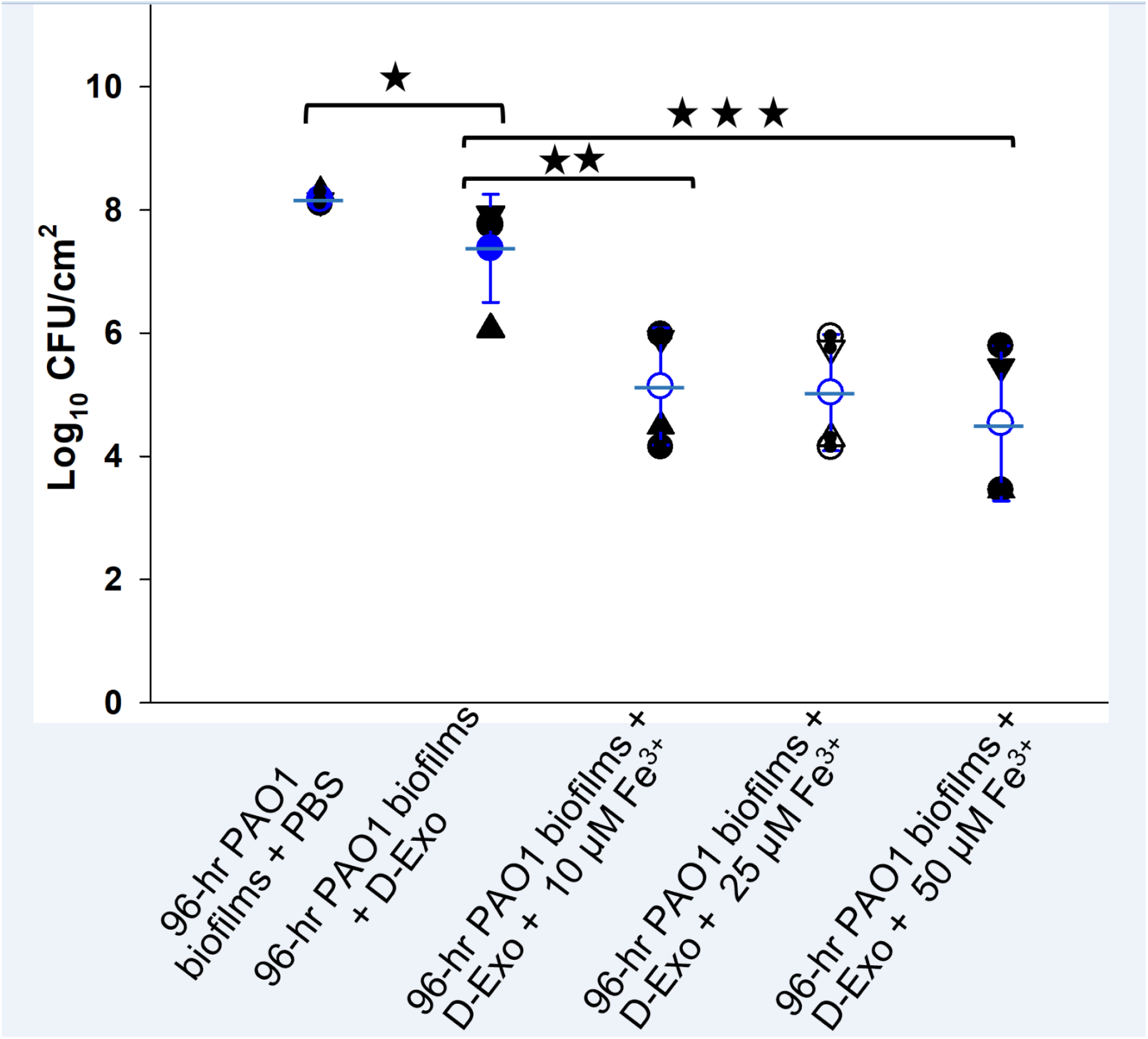
Synergic effects of D-Exo/Fe^3+^ on 96-h *P. aeruginosa* PAO1 biofilm growth. For all date points, biomasses of biofilms were collected and analyzed 24 hours after each treatment. The mean value and error bar of each group of data are given in blue. P-value <0.05 (*), <0.03 (**) and <0.01(***).

Using a commercial cellular ROS assay kit, we observed that the biofilms treated with D-Exo produced much higher levels of ROS than the negative control biofilm (treated with PBS buffer) and the positive control biofilm (treated with tetracycline at 10 mg/mL) (**Fig. 5**). Although specific ROS species were not identified in this measurement, these results provide evidence that D-Exo promotes iron uptake and induces overall ROS accumulation within the biofilm, ultimately leading to cell death.

**Fig. 5.**
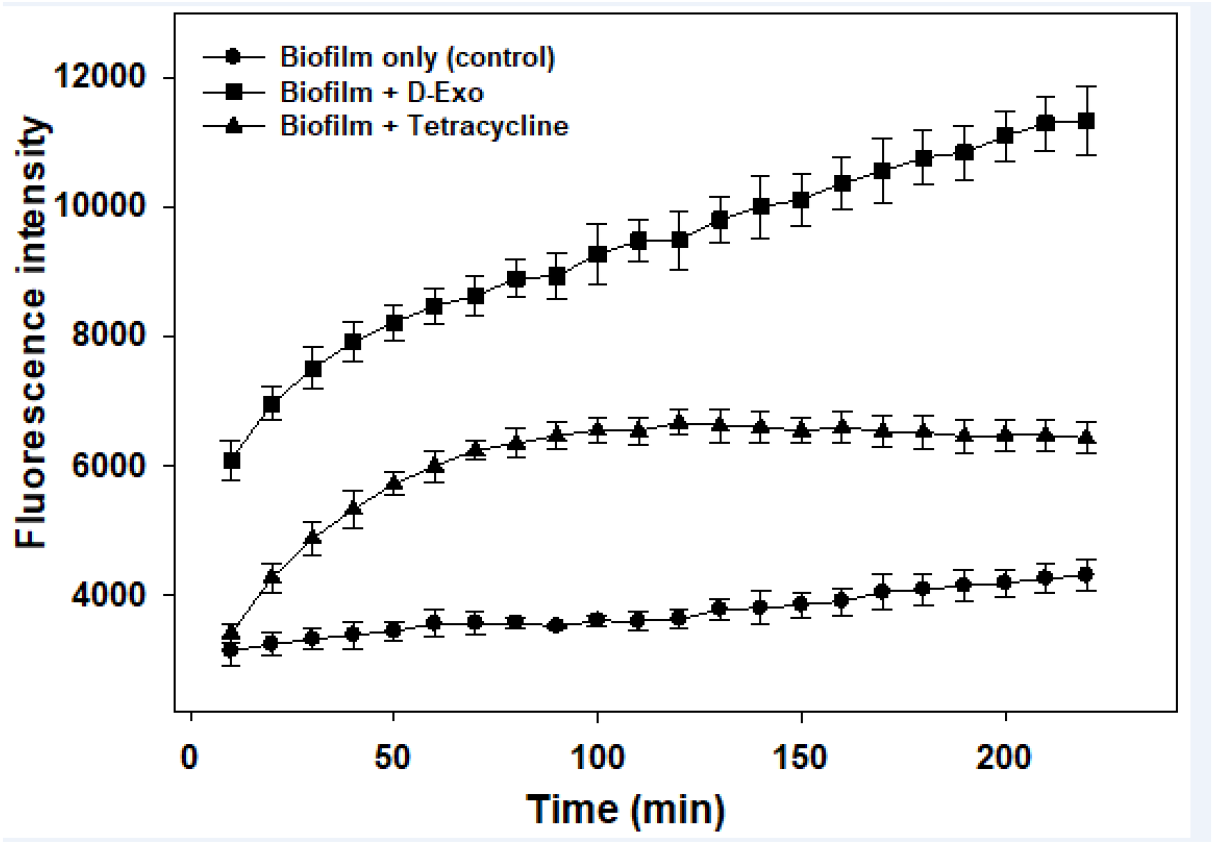
ROS levels of *P. aeruginosa* PAO1 biofilms in the lag phase were monitored under various conditions. After 1 hour of incubation with dye agents from a Cellular ROS Assay Kit (Red) (ab186027), *P. aeruginosa* PAO1 bacteria in the lag phase were mixed with PBS buffer, tetracycline (10 mg/ml), or D-Exo (97.9 μg/ml), immediately followed by monitoring of changes in their fluorescence intensity (excitation/emission: 520/605 nm). An increase in fluorescence intensity was correlated with an increase of overall ROS levels in each sample.

To determine the potential cytotoxicity of *P. aeruginosa* PAO1 D-Exo on mammalian cellular function, we examined the viability of human mesenchymal stem cells (hMSC) in the presence of the D-Exo (0.33 μg/μL) (see *<u>Supplementary Information section S5</u>*). The results showed no significant detrimental effects on either cell growth (**Fig. S7A**) or cellular morphology (**Fig. S7B**) of hMSC after exposure to the D-Exo for two days. These observations are consistent with literature studies on the biosafety of using bacterial exosomes in human bone and tissue therapy [39].

## Discussion

Bacterial biofilms are complex surface-attached communities of bacteria held together by self-produced polymer matrices. The formation of biofilms is a complex process and involves multiple stages, or phases. As they enter each of these development phases, bacteria quickly respond to changes in the environment to propagate or survive by rapidly communicating with neighboring cells and reorganizing their intracellular physiological processes. Bacterial exosomes are believed to be key players in bacterial intercellular communications.

In this study, we are the first to report that exosomes secreted by *P. aeruginosa* PAO1 biofilms under different environmental conditions can have different effects on the same population of *P. aeruginosa* PAO1 biofilms, meaning that the functional effects of exosomes are conditional, or environment-dependent. For example, the growth of 2-h *P. aeruginosa* PAO1 biofilms can be enhanced by 2-log_10_ CFU/cm^2^ in the presence of their G-Exo (**Fig. 2A**), while the presence of their D-Exo can eliminate 8-h and 24-h *P. aeruginosa* PAO1 biofilms (**Fig. 2B** and **Fig. 2C**). These results suggest that the functional roles of exosomes are closely correlated with the functions of the parental cells under the specific conditions under which the exosomes are secreted.

Because of the rich nutrition and favorable growth conditions of the exponential growth phase, bacteria optimize their internal cellular processes to expand their population rapidly and establish biofilm communities by activating bacterial attachments [40]. During this active growth period, bacteria likely secrete G-Exo as messengers to coordinate the cellular actions of neighbors to establish biofilm communities. The proteomic analysis of G-Exo and biomass from the exponential growth phase identified multiple surface proteins that may play a role in promoting bacterial attachment and biofilm development (**Table 1** and **Table S1**).

Once bacterial biofilm is established, the community enters the stationary plateau phase, in which the death and growth rates are balanced and the biofilm matures. At this stage, the bacteria within the biofilm become more resistant to antimicrobial treatments [41]. Under normal culture conditions or in favorable growth environments, mature biofilms ultimately enter the dispersal/detachment phase, during which bacterial cells depart from the biofilm to start or participate in a new biofilm formation cycle. Because of the decrease in available nutrients, cells start to starve. The starvation affects the biofilm in two ways. First, some of the bacterial cells within the biofilm die to generate nutrients by decreasing the population density (increasing the amount of nutrients per cell) and by releasing nutrients through dead cells to provide nutrition for the remaining nutrient-deprived cell population [34, 42-46]. Second, to survive in such an extreme condition, the core group of biofilm bacteria rapidly adjusts its intracellular metabolic pathways, leading to alterations in intracellular components and surface proteins. Meanwhile, the exosomes (the D-Exo) are secreted by the core bacteria to reflect these alterations. The secreted D-Exo then serve as cellular messengers to coordinate the actions of the core population of bacteria and enhance their ability to acquire vital supplies and nutrition from a resource-scarce environment.

Several unique proteins were identified on the surface of the D-Exo (**Table 1**), all belonging to a group of proteins that aid bacterial growth by acquiring iron resources from nutrition-depleted environments. Iron is an essential component of metabolic enzymes and regulatory proteins; it acts as an electron carrier and a key nutrient for bacterial life and thus plays a critical role in bacterial physiology to support growth. Therefore, it is likely that D-Exo serves as a unique messenger to help bacterial survival under conditions of nutrition/iron deprivation by coordinating bacterial cellular functions and delivering key proteins to the recipient cells to enhance their nutrition acquisitions. However, D-Exo can only benefit recipient bacterial cell growth/survival under nutrition depletion conditions. In normal or iron-rich culture environments, D-Exo becomes toxic to bacteria. For instance, as shown in **Fig. 2**, the presence of D-Exo can effectively inhibit the growth of *P. aeruginosa* PAO1 biofilms under normal culture conditions. Furthermore, the inhibition power of D-Exo against 96-h *P. aeruginosa* PAO1 biofilms can be significantly enhanced by the presence of extra Fe^3+^ (**Fig. 4**).

Maintaining an intracellular iron concentration of 10^-6^ M is essential for bacterial growth; therefore, iron uptake is strictly regulated by bacterial cellular processes [31, 47]. Insufficient iron uptake can lead directly to cell starvation, while excessive iron uptake in a medium-rich environment can result in bacterial cell death by activating ferroptosis, an iron-dependent form of programmed cell death [48, 49]. A consequence of ferroptosis activated by a high intracellular concentration of iron is the triggering of the Fenton/Haber-Weiss reaction, which produces lethal concentrations of ROS, leading to cell death [31]. This is supported by the results shown in **Fig. 5**, which demonstrate that *P. aeruginosa* PAO1 bacteria produced more overall ROS when exposed to D-Exo than when exposed to an antibiotic drug. These findings are significant because they provide a foundation for utilizing bacterial D-Exo as a novel antibiotic alternative to address the bacterial biofilm-associated drug-resistant infections that currently threaten global human health.

In summary, our study on the growth of *P. aeruginosa* PAO1 bacterial biofilms in response to the exosomes secreted by the biofilms provides strong evidence of the importance of bacterial exosomes in regulating biofilm growth. We are the first to show that the regulatory roles of bacterial exosomes depend on the biofilm developmental stage at which the exosomes are secreted. In addition, we discovered dual roles of D-Exo in controlling biofilm growth: 1) help bacterial survival under nutrition depletion conditions and 2) treat *P. aeruginosa* PAO1 biofilms in the presence of ferric ions. These significant discoveries bring both excitement and challenge to the field. The excitement is that the discoveries, especially the Fe^3+^-enhanced inhibition power of D-Exo, pave the way to developing new therapies against drug-resistant bacterial infections. The challenge is that our results raise fundamental questions. For example, how do environmental signals/conditions at each stage of bacterial biofilm development affect exosome biogenesis? Do exosomes secreted by other types of bacteria have the same properties and use a similar mechanism to inhibit biofilm growth? Are there other mechanisms involved? What are the roles of exosomal DNA/RNA components in the mechanisms, and how do they change when exosomes are secreted by bacteria at different developmental stages? How do intracellular pathways of biomass change in response to exposure to D-Exo? Answers to these questions will shed light on the regulatory roles and detailed mechanisms of bacterial exosomes in biofilm growth and how the roles switch from one stage to another. Furthermore, our data suggest that the inhibition of biofilm growth by D-Exo involves a large amount of ROS production. However, the nature of the ROS species involved in the inhibition is unknown. Answering this question will help us pinpoint the intracellular mechanism of D-Exo-induced cell death. Moreover, is it possible that the observed condition-dependent regulatory roles of bacterial exosomes can be applied to other cellular systems? Answering this question may help address the challenges faced in the clinical application of mammalian or stem-cell-derived exosome-based therapies for various disease treatments [50-52].

## Materials and methods

### 1.1 Biofilms

*P. aeruginosa* (PAO1) [53] from a -80°C freezer was streaked on tryptic soy agar (TSA, BD Difco™ Dehydrated Culture Media, DF0369-17-6) and incubated overnight at 37°C. The *P. aeruginosa* was maintained on TSA plates at 4°C and an inoculum was prepared by transferring a touch from this plate into 10 mL of tryptic soy broth (BD Bacto™) overnight. Then, a volume of 100 μL (OD_600_=0.1) was used to seed 47-mm sterilized filter membrane (Membrane Solutions MCE Gridded Membrane Filter, Mixed Cellulose Esters Membrane Filter, Pore Size: 0.45 μm) on TSA petri dishes. The plates were incubated at either 37.0°C or 32.7°C. At desired time points (0, 1, 2, 4, 8, 12, 16, 24, 32, 48, 72, 85, 96, and 120 h), the biofilm was suspended in 0.9% NaCl to prepare serial dilutions for colony forming unit (CFU) counting. The viable cell concentration (CFU/cm^2^) was calculated according to the literature [54, 55]. All experiments were done under aseptic conditions. TSA and TSB were autoclaved at 121°C and 1.5 psa for 50 min/L.

### 1.2 Exosome extraction

Biofilms for exosome extraction were grown at 32.7°C on 47-mm sterilized membrane paper (Membrane Solutions MCE Gridded Membrane Filter, Mixed Cellulose Esters Membrane Filter, Pore Size: 0.45 μm) on TSA petri dishes for 8 hours or 96 hours. Exosomes were isolated and purified from *P. aeruginosa* PAO1 biofilms using a conventional differentiation centrifugation protocol [21-24]. Briefly, an inoculum with 4.4x10^5^ ± 7.3 x10^2^ CFU/cm^2^ was used to seed 47-mm sterilized membrane paper on TSA plates and incubated. When biofilms reached their exponential growth phase their exosomes were extracted and defined as growth exosome (G-Exo). Similarly, when biofilms reached their death/survival growth phase, their exosomes were extracted and defined as death exosome (D-Exo). These processes are illustrated in **Fig. S1**. For exosome extraction, the biofilms were removed from the membrane and then were suspended in TSB. The cell debris was removed by centrifugation at 10,000 g for 30 min at 4°C using a benchtop centrifuge (Biofuge, HERAEUS). Then, the supernatant was collected and filtered using a 0.45-μm syringe filter (GenClone 25-246, Syringe Filters, PES, 30-mm Diameter, Sterile, Cat #: 25-246). The filtrate was ultracentrifuged (Optima LE-80K Ultracentrifuge, Beckman) at 150,000 g for 2 hours at 4°C, twice, and filtered with a 0.22-μm filter (GenClone 25-244, Syringe Filters, PES, 30-mm Diameter, Sterile, Cat #: 25-244) after the first run to remove macrovesicles >200 μm in size. PBS was used as a buffer to suspend the pellets after each ultracentrifugation run. The extracted exosomes were stored at -20°C for further analysis.

### 1.3 Testing activity of exosomes against biofilm at different ages

To investigate the functional effect of extracted exosomes on the growth of *P. aeruginosa* PAO1 biofilms, we used the Kirby-Bauer disk diffusion susceptibility test [56, 57]. Briefly, an inoculum was prepared by activating a touch from the *P. aeruginosa* PAO1-TSA plates in TSB overnight. Then, the bacterial suspension was adjusted to OD600 ∼0.1, and the adjusted suspension was spread onto TSA plates using 6-inch sterilized cotton swabs and left for 2 hours. Discs were loaded with 25 μL of PBS; tetracycline (Cat # 87128, Sigma-Aldrich) at 0.033 μg/μL, 0.22 μg/μL, and 1 μg/μL; G-Exo at 0.037 μg/μL; and D-Exo at 0.11 μg/μL, 0.22 μg/μL, and 0.33 μg/μL, respectively. The treated discs were then placed on the bacteria-seeded TSA plates and incubated at 37°C overnight.

To quantify the effects of G-Exo and D-Exo on the growth behavior of *P. aeruginosa* PAO1 biofilms, biofilms grown for 2 h (lag phase), 8 h (exponential phase), and 24 h (stationary phase) were tested. Briefly, an inoculum was prepared by transferring a touch from the *P. aeruginosa* PAO1-TSA plates into 10 ml of TSB and incubating the plates at 37°C overnight. A volume of 2.5 μL (OD_600_ ∼0.5) was used to seed sterile 13-mm polycarbonate membrane (WHA10417401, Sigma-Aldrich) on TSA plates and incubated at 37°C. Biofilms grown for 2 h were treated with one dose of G-Exo (0.028 μg/μL) or D-Exo (0.33 μg/μL) and incubated at 37°C for 24 hours. Biofilms grown for 8 h were treated with one dose of G-Exo (0.037 μg/μL) or D-Exo (0.33 μg/μL) and incubated at 37°C for 24 hours. Three doses of D-Exo (0.33 μg/μL) were applied (one dose every 24 h) to biofilms grown for 24 h, and the plates were incubated at 37°C. Then the viable cells were counted and reported as CFU/cm^2^.

### 1.4. Bacterial cells, biofilms and exosome imaging

We used a Leica SP-5 confocal laser scanning microscope to image biofilms grown for 8 h. The biofilms, planktonic cells and controls were stained with Bacteria LIVE/DEAD BacLight Bacterial Viability Kits following published literature[58]. Excitations/emissions of 480/500 nm for SYTO 9 and 490/635 nm for propidium iodide were used.

Transmission electron microscopy (TEM, FEI Tecnai G2 20 Twin equipped with a 200-KV LaB6 electron source) was used to image single cells and exosomes. *P. aeruginosa* PAO1 biofilms grown for 8 h were incubated with D-Exo for 24 hours. Cells were suspended in 1% PBS for 5 minutes, and then 3 drops of 1% PBS and one drop of osmium were added and incubated at -4°C overnight. After the resin was removed, the samples were washed with distilled water and exposed to a series of ethanol concentrations (30%, 50%, 70%, 90%, and 100%). Subsequently, the cells were mixed with Spurr in propylene oxide and left on the shaker overnight. The next day, 100% Spurr was added, and the samples were kept at 60°C for 24 hours. Blocks were trimmed, and 70-nm thin sections were prepared and loaded onto formvar/carbon-coated copper EM grids with a thickness of 200 nm. The grids were positively stained with 2% uranyl acetate and lead. Grids were examined using TEM.

The direct interactions between G-Exo and bacterial cells were investigated using TEM and CLSM. To investigate using TEM, 8-h *P. aeruginosa* PAO1 biofilms were incubated with G-Exo for 24 hours. Four μL of the cell sample were loaded onto formvar/carbon-coated copper EM grids with a thickness of 200 nm. The grids were then stained with 4 μL of 2% uranyl acetate and examined using TEM. For CLSM, 2-h *P. aeruginosa* PAO1 biofilms were incubated with 10 μL of DAPI-dye-labelled G-Exo at 37°C in the dark for 24 hours. DAPI (diamidino-2-phenylindole) stains exosomal DNA. Five μL of the treated cell suspension were loaded onto a glass slide and examined with CLSM using an excitation/emission wavelength of 359/461 nm.

### 1.5 Qualitative fluorescence-based assay of exosomes

Exosomes were labeled with Vybrant™ DiI Cell-Labeling Solution according to the manufacturer’s protocol. To prepare the staining medium, 5 μL of the labeling solution is added to 1 mL of PBS. The exosomes are then incubated with the staining medium at 37°C for 20 minutes. After incubation, a small volume (5 μL) of the stained suspension is placed on a glass slide and examined under a confocal laser scanning microscope (Leica SP-5 Confocal Laser Scanning Microscope). The excitation and emission wavelengths used for imaging are 549 and 565 nm, respectively. This method can provide valuable insights into the cellular uptake and distribution of exosomes labeled with the fluorescent dye [59-61].

### 1.6 Quantitative protein-based analysis of exosomes

To quantify the protein concentration of the extracted exosome samples, a modified bicinchoninic acid assay (BCA) protocol was used (Pierce™ BCA Protein Assay Kit, Thermo Fisher Scientific) [62]. The protocol involved denaturing equal amounts of extracted exosomes (25 μL) using RIPA 5x for 30 min on ice, followed by protein precipitation after the addition of trichloroacetic acid (TCA, Sigma-Aldrich, 91228), (v:v, 1:0.25 of sample: TCA). The resulting precipitate was collected by centrifugation at 6000 rpm for 5 min and washed with cold acetone. The precipitate was then dissolved in PBS (25 μL) and mixed with an equal amount of the working solution (50 parts reagent A, 48 parts reagent B, and 2 parts reagent C) in a 96-well plate. The plate was incubated at 37°C for 2 hours, and the absorbance was measured at 562 nm using a plate reader (Cytation 5 imaging reader).

### 1.7 Proteomic analysis

Proteomic identifications of extracted G-Exo and D-Exo were done using proteome discoverer (Thermo Orbitrap Fusion Tribrid). A 100-μL volume of exosomes was centrifuged for 1 h at 4°C and 150,000xg. The pellet was dissolved in 50 μL of 6 M guanidine hydrochloride (GuHCL, Thermo Scientific™, AAJ6078622) and placed on the shaker for 20 min at 1000 rpm; then, samples were sonicated 4 times for 10 s with a 2-min ice cooling interval and stored at -70 °C.

### 1.8 Reactive oxygen species detection

We used a Cellular ROS Assay Kit (Red) (ab186027) to measure reactive oxygen species (ROS) in our biofilm samples following the manufacturer’s protocol (Abcam). A 25-μL (OD600=0.1) sample was transferred to each well in a 96-well plate and incubated for 7 h. A 100-μL volume of ROS working solution was added to each well and incubated for 1 h, followed by the addition of D-Exo (0.098 μg/μL), tetracycline (10 mg/ml), or PBS. The plate was placed in the plate reader for immediate monitoring of the change in ROS levels by measuring the fluorescence increase at excitation/emission 520/605 nm (cutoff 590 nm). The ROS working solution was prepared by mixing 40 μL of DMSO with the red stain; then 5 μL of the mixture was mixed with 2.5 ml of buffer.

### 1.9 Statistical analysis

All experiments were done in biological replicates, and the number of biological replicates is given for each experiment. The data were averaged and are presented as the standard error of the mean. The statistical significance of the differences between treatments was determined using the Wilcoxon rank sum test. A p-value less than 0.05 was considered significant [63].

## Acknowledgements

This study is partially supported by NIH (R21CA248734) to WJD. MGS was primarily supported by the graduate school of Washington State University. We thank Mr. Md Monzurul Islam Anoy for his help in reviewing protocols of biofilm data collection and analysis.

## Author contributions

MGS worked out most experimental details, conducted the majority of the experiments and data analysis, and contributed to the manuscript writing. HB contributed to the design of the reactive oxygen species investigation and proteomic analysis, provided partial budget support for the project, and contributed to manuscript revisions. WJD developed the project, conceptualized the main ideas and hypotheses, developed strategies for testing the hypotheses, provided budget support, and was responsible for finalizing the manuscript.

All the authors approved the final version of the manuscript.

We declare no conflicts of interest.

