## Supplementary material for "Dual Roles of the Conditional Exosomes Derived from *Pseudomonas aeruginosa* Biofilms: Promoting and Inhibiting Bacterial Biofilm Growth": Fig. S1

**Classifications**

Major category: Biological Sciences

Minor category: Microbiology

**Keywords:**

Bacterial exosomes, biofilm, *Pseudomonas aeruginosa,* biofilm control

#### S1. Exosome extraction

PAO1 bacterial inoculum with $4.4 \times 10^5 \pm 7.3 \times 10^2$ CFU/cm$^2$ was used to seed a 47-mm sterilized membrane on sterilized TSA plates and incubated in the dark at 32.7° C. When the bacterial culture reached the desired growth phase (as shown in **Fig. 1**), G-Exo and D-Exo were isolated and purified following the procedure shown in **Fig. S1.**

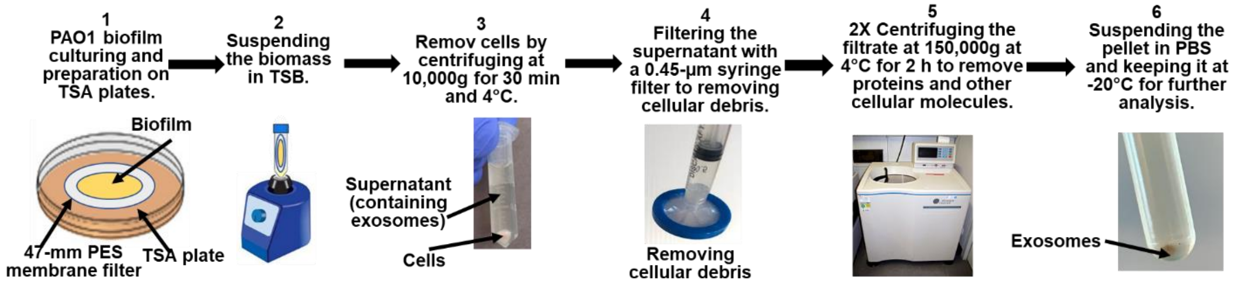


**Fig. S1.** The exosome extraction protocol consisted of 1) culturing PAO1 biofilm on filter membrane, 2) suspending biofilm cells to release exosomes, 3) spinning down the bacterial cells, 4) filtering the supernatant through a 0.45-µm filter to remove cellular debris, 5) centrifuging twice at ultrahigh speed to separate exosomes from soluble proteins and other cellular molecules, and 6) suspending the exosome pellets in PBS buffer and keeping the extracted exosomes at -20ºC for further analysis.

#### S2. Correlating the protein concentration of exosomes to the particle numbers

To correlate the protein concentration of G-exosomes to their particle numbers, purified exosomes were quantified by their content protein concentration (µg/ml) using a Pierce™ BCA Protein Assay Kit and stained with Vybrant™ DiI Cell-Labeling Solution (see the Materials and Methods section). The number of fluorescent exosome particles in each image (**Fig. S2**) was correlated to their protein concentration. The protein concentration is plotted against the exosome particle number in **Fig. S3**, which shows a linear relationship between them.

$$Y(\text{number}*10^3) = 24450P(\mu g/ml) - 458385; \qquad (R^2 = 0.979)$$

where Y is the exosome particle number and P is the exosomal protein concentration (µg/ml). Based on this relationship, the average protein content of each exosome particle is 0.00188

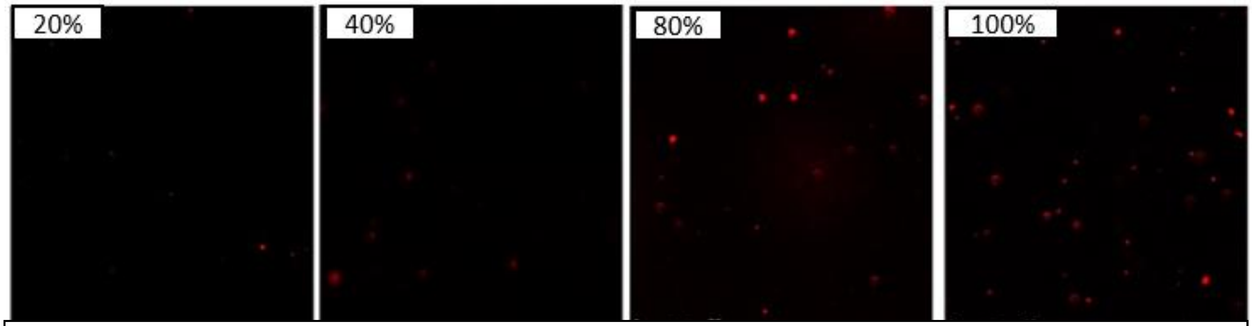


**Fig. S2. Quantitative fluorescence-based assay of exosomes.** Exosomes stained with Vybrant™ DiI Cell-Labeling Solution (red spots) were imaged under a confocal fluorescence microscope with an excitation/emission of 549/565 nm. The pictures from left to right show exosomal protein concentrations from low to high (20%= 26.4 ± 3 µg /ml, 40%= 40.4 ± 1 µg/ml, 80%= 78.7 ± 1 µg/ml, and 100%= 96.4 ± 4µg/ml). The scale bar indicates 75 µm.

$\pm 8 \times 10^{-4}$ µg/ml.

#### S3. Shape and size of exosomes

Ten microliters of each exosome sample were diluted with PBS at a v/v ratio of 1:1000. Four microliters of the diluted sample were then loaded onto formvar/carbon-coated copper EM grids with a thickness of 200 nm. After drying, the grids were stained with 2% uranyl acetate (4 µl) and the excess stain was immediately removed. The grids were then allowed to dry under light for 15 minutes. Finally, the grids were examined using a transmission electron microscope (FEI Tecnai G2 20 Twin equipped with a 200KV LaB6 electron source). TEM images of the purified G-Exo and D-Exo are shown in **Fig. S4**. The size of exosomes extracted during the exponential growth phase was larger (112.9 ± 3.7 nm) than that of exosomes extracted during the death/survival phase (33.16 ± 0.86 nm).

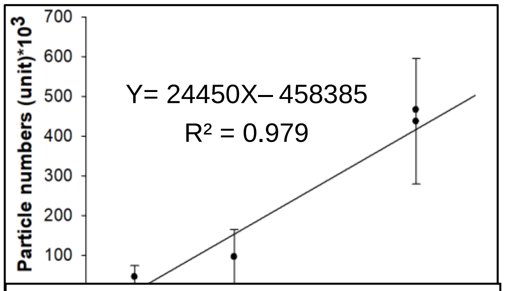


**Fig. S3.** Plot of the linear relationship of exosomal protein concentration (µg/ml) to exosome particle number (x$10^3$). Data points indicate the mean, and error bars indicate standard deviations (n = 8).

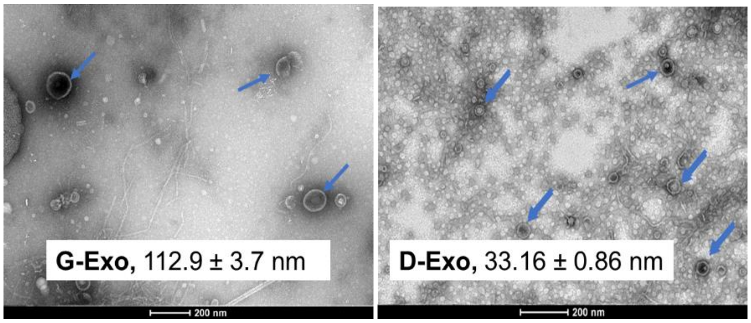


**Fig. S4.** TEM images showing the sizes and shapes of the G-Exo and D-Exo from PAO1 biofilms. Samples are negatively stained with 2% uranyl acetate. Magnification power = 25000X.

#### S4. Functional effects and importance of intact exosome structure of bacterial exosomes

To investigate the functional effects of D-Exo PAO1 biofilm growth, we conducted a disc diffusion assay (see the Materials and Methods section) using both D-Exo and G-Exo to observe their functional effects on the growth behavior of PAO1 biofilm (**Fig. S5A**). PBS buffer and the antibiotic tetracycline were used as negative and positive controls, respectively, to assess the efficacy of D-Exo/G-Exo in affecting the growth of PAO1 biofilms. Given that PAO1 is a known antibiotic-resistant superbug, we used a final concentration of tetracycline of 1 µg/µl in the test to make it comparable to other treatments. G-Exo did not show an inhibition/clear zone, and D-Exo exhibited effective and dose-dependent inhibition of PAO1 biofilm growth across a range of exosomal protein concentrations (0.11, 0.22, and 0.33 µg/µl).

Although we believe that D-Exo-mediated cellular communications are key to inhibiting biofilm growth, it is possible that the observed effects are caused by toxins associated with D-Exo, rather than by D-Exo-based cellular communications. To rule out toxin-based effects and confirm that the intact exosome structure is critical for inhibiting bacterial biofilm growth, we performed a diffusion susceptibility test to assess the inhibition effects of lysed D-Exo. In this test, the cell walls of the exosomes were lysed by subjecting them to three 10-second water bath sonication cycles, with a 2-minute interval in ice. The results (see

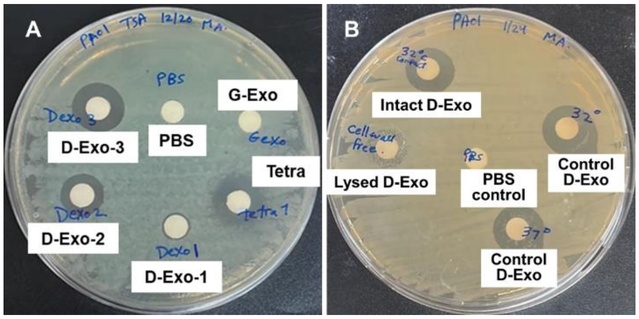


**Fig. S5. A:** Kirby-Bauer disk diffusion susceptibility test of the effects of G-Exo (36.9 µg/ml) and D-Exo (1, 2, and 3 indicate 0.11, 0.22, and 0.33 µg/µl, respectively) on PAO1 biofilm growth. Tetracycline and PBS buffer were used as a positive control and a negative control, respectively. Note: to generate a visible clear control zone, 1 µg/µl of tetracycline was used, which is much higher than the medical tolerated doses (0.033-0.22 µg/µl) used for adult admissions [1]. **B:** Diffusion susceptibility test of the effect of the intact D-Exo (0.33 µg/µl) and lysed D-Exo (0.33 µg/µl) on PAO1 biofilm growth. The lysed D-Exo showed much less inhibition of biofilm growth than the intact D-Exo.

**Fig. S5B**) showed a clear zone of biofilm inhibition around the filter disc containing intact D-Exo, whereas the zone around the disc containing lysed D-Exo was much fainter. These results suggest that the intact exosome structure, rather than toxin components, plays a role in regulating biofilm growth.

#### S5. Mechanistic investigations of inhibition effects of the D-Exo on biofilm growth

To test the hypothesis that the suppression of PAO1 biofilm growth is caused by D-Exo-induced excessive iron uptake by D-Exo recipient cells leading to the activation of bacterial cell death, we examined whether D-Exo-induced excessive iron uptake could improve the poor efficacy of the D-Exo against 96-h biofilms (**Fig. 2D**). Briefly, we first treated 96-h PAO1 biofilms with 2 doses of D-Exo at a protein concentration of 0.33 µg/µL with a 12-hour interval, which only induced an inhibition of less than 1-$log_{10}$ (**Fig. S6A**). We then repeated the same inhibition experiments applying 10 µM, 25µ M, or 50 µM of $Fe^{3+}$ directly to the 96-h PAO1 biofilms to see how iron alone affects the growth of 96-h PAO1 biofilms. Compared to the PBS control, the presence of ferric ions alone marginally promoted biofilm growth with a dose-dependent effect.

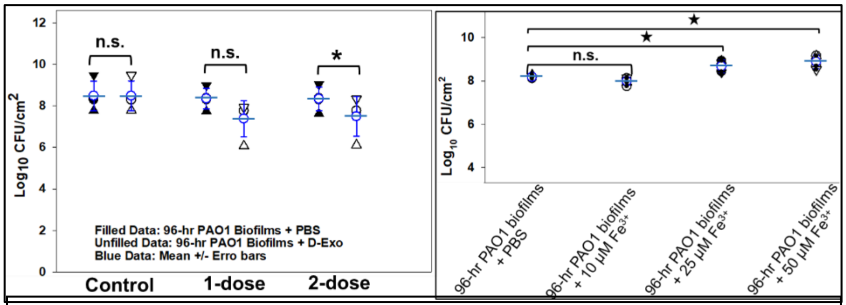


**Fig. S6**. **A:** Inhibition effects of D-Exo on 96-h PAO1 biofilms. Two doses of D-Exo at a protein concentration of 0.33 µg/µL were applied to the biofilms with a 12-hour interval, which induced an inhibition of less than 1-$log_{10}$. **B:** Effects of 10 µM, 25 µM and 50 µM of ferric ions on 96-h biofilm. For all data points, biomasses of biofilms were collected and analyzed 24 hours after each treatment. Ferric ions alone showed a very minimal effect on the biofilm growth. The mean value and error bar of each group of data are given in blue. P-value <0.05 (*). n.s.: not significant.

#### S6. Cell viability of D-Exo derived from PAO1 biofilms

We investigated the toxicity of PAO1 D-Exo (0.33 µg/µl) to mammalian cell viability. Human mesenchymal stem cells (hMSC) obtained from ATCC (Manassas, VA) were cultured in alpha-MEM medium with 5% FBS, 1% L-glutamine, and 1% Pen-strep in an incubator with 5% $CO_2$ at 37°C. Cells were then used to seed 6-well plates with an inoculation rate of 10^4 and cultured to 60-70% confluency before being exposed to D-Exo (0.33 µg/µl) or PBS (as a control) in separate cultures of hMSC. After 24 hours and 48 hours of exposure, the growth and cellular morphologies of the hMSC were examined. The results, depicted in **Fig. S7A** and **Fig. S7B**, showed no

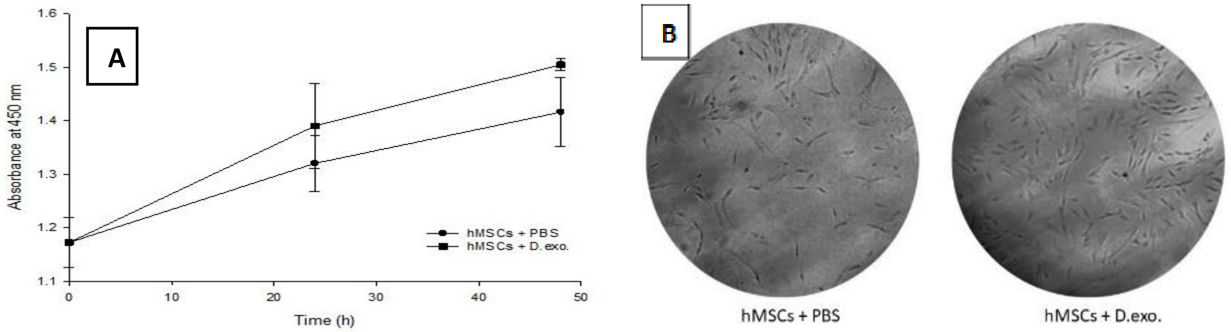


**Fig. S7. Cytotoxicity test of D-Exo to human mesenchymal stem cell culture. A.** Changes in hMSC growth in the absence/presence of D-Exo (0.33 µg/µl) were monitored for two days. At each time point, cells were stained with Cell Proliferation Reagent WST-1 and incubated for four hours. The absorbance was measured at 450 nm to monitor the replication of their genomic DNA. **B.** Light microscopy images showing changes in cellular morphology of hMSCs 48 hours after treatment with D-Exo (0.33 µg/µl).

significant detrimental effects on either cell growth or the cellular morphology of hMSC after two days of exposure to D-Exo. These results demonstrate that the presence of D-Exo does not affect cellular functions and thus provide a foundation for exploring the use of D-Exo in antibiotic applications.

### Table S1. Summary of proteomic analysis of G-Exo

| Protei | Accession | Description | Exp. q | Sum PEF | Coverag | # PSMs ( | # Unique | # AAs | MW [kDa | Abundanc |
|---|---|---|---|---|---|---|---|---|---|---|
| 1 | 2 | 3 | 4 | 5 | 6 | 7 | 8 | 9 | 10 | 11 |
| Growth promotors | | | | | | | | | | |
| High | A0A653B731 | Cell division proteins | 0 | 6.093 | 4 | 1 | 1 | 398 | 41.7 | 100 |
| High | V6A9F4 | Phasin_2 domain-containing protein | 0.004 | 1.275 | 12 | 2 | 1 | 153 | 16.9 | 34.64 |
| High | A0A1C7B9B8 | Elongation factors | 0 | 48.011 | 23 | 22 | 11 | 706 | 77.7 | 58.34 |
| High | A0A7W3UUK | Chaperonins | 0 | 11.74 | 10 | 7 | 1 | 527 | 55.7 | 32.5 |
| High | A0A3D9EJG5 | Chaperone(s) | 0.002 | 1.443 | 4 | 1 | 1 | 433 | 47.9 | 35.28 |
| DNA synthesis | | | | | | | | | | |
| High | A0A072ZFF8 | N5-carboxyaminoimidazole ribonucleotide mutase | 0 | 37.121 | 67 | 22 | 5 | 163 | 16.9 | 60.32 |
| High | A0A086BUQ2 | Vitamin B12-dependent ribonucleotide reductase | 0 | 14.186 | 7 | 4 | 3 | 734 | 82.7 | 64.4 |
| High | S6ABT0 | Ribonucleotide reductase | 0 | 3.464 | 3 | 3 | 1 | 324 | 35.6 | 33.98 |
| High | A0A3M5EFC( | Polyribonucleotide nucleotidyltransferase | 0 | 116.971 | 34 | 60 | 16 | 748 | 80.3 | 49.36 |
| High | A0A367M4C8 | HU family DNA-binding protein | 0 | 27.865 | 49 | 19 | 5 | 93 | 9.8 | 68.58 |
| High | A0A069PX19 | DNA polymerases | 0 | 18.819 | 11 | 7 | 6 | 913 | 99.7 | 61.2 |
| High | A0A086BZZ8 | DNA topoisomerase | 0 | 13.875 | 10 | 6 | 5 | 868 | 97.2 | 56.28 |
| High | Q9I3X2 | DNA helicase | 0 | 7.812 | 6 | 3 | 3 | 711 | 79.8 | 41.14 |
| High | A0A485FIC3 | DNA gyrase | 0 | 41.307 | 11 | 18 | 6 | 925 | 101.3 | 43 |
| High | A0A2S5IJE8 | Holliday junction ATP-dependent DNA helicase | 0.002 | 1.637 | 6 | 1 | 1 | 205 | 22.3 | 69.12 |
| Protein processing | | | | | | | | | | |
| High | A0A086C2F0 | Peptide-binding protein | 0 | 9.811 | 18 | 3 | 3 | 222 | 24.1 | 100 |
| High | A0A397MC94 | SsrA-binding protein | 0.001 | 2.281 | 6 | 1 | 1 | 197 | 22.3 | 100 |
| High | A4XZJ8 | Putative serine protein kinase | 0 | 14.827 | 11 | 9 | 1 | 640 | 73.7 | 100 |
| High | A0A485EER1 | NAD-capped RNA hydrolase | 0 | 3.84 | 2 | 1 | 1 | 841 | 92.3 | 100 |
| High | A0A367M033 | 2-oxo-4-hydroxy-4-carboxy-5-ureidoimidazoline decarboxylas | 0 | 3.647 | 9 | 1 | 1 | 178 | 19.6 | 100 |
| High | A0A080VRC4 | S-adenosylmethionine:tRNA ribosyltransferase-isomerase | 0 | 3.558 | 4 | 1 | 1 | 347 | 38.1 | 100 |
| High | A0A5E9K219 | Ribosomal RNA large subunit methyltransferase H | 0.001 | 2.577 | 12 | 2 | 1 | 155 | 17.8 | 100 |
| High | A0A072ZEM5 | Aminopeptidase P family protein | 0.001 | 2.244 | 3 | 1 | 1 | 405 | 44.1 | 100 |
| High | A0A0H2ZCX7 | Putative MoxR protein O | 0.001 | 2.446 | 4 | 1 | 1 | 305 | 32.8 | 100 |
| High | A0A1I1XAM6 | Adenylyltransferase and sulfurtransferase | 0.002 | 1.941 | 4 | 1 | 1 | 270 | 28.4 | 100 |
| High | W1MVH1 | Glutathione S-transferase | 0.004 | 1.38 | 5 | 1 | 1 | 256 | 28.8 | 100 |
| High | A0A0A8RCC8 | Periplasmic tail-specific protease | 0.001 | 2.185 | 1 | 1 | 1 | 840 | 93.5 | 100 |
| High | A0A072ZJB8 | Carboxy-terminal processing protease precurs | 0.001 | 2.716 | 8 | 2 | 2 | 436 | 46 | 31.7 |
| High | B7V669 | 50S ribosomal proteins | 0 | 56.677 | 49 | 26 | 7 | 129 | 14.5 | 42.56 |
| High | A0A1C7C943 | 30S ribosomal proteins | 0 | 35.017 | 28 | 26 | 4 | 246 | 27.3 | 33.62 |
| High | A0A231K3D3 | Chain-length determining protein | 0 | 16.217 | 17 | 5 | 4 | 442 | 49.2 | 8.08 |
| High | W1MWV0 | Glycine--tRNA ligase alpha subunit | 0 | 13.843 | 24 | 7 | 5 | 318 | 36.5 | 42.18 |
| High | A0A0C7D284 | Tyrosine--tRNA ligase | 0 | 6.676 | 10 | 2 | 2 | 399 | 44.1 | 80.42 |
| High | A0A0A8RE97 | Leucine--tRNA ligase | 0 | 6.149 | 3 | 4 | 2 | 873 | 97.6 | 30.8 |
| High | Q9HXU0 | Lysine--tRNA ligase | 0 | 11.043 | 12 | 4 | 4 | 501 | 57.3 | 72.02 |
| High | A0A0F6RS41 | Phenylalanine--tRNA ligase | 0.004 | 1.284 | 2 | 1 | 1 | 792 | 86.7 | 61.18 |
| High | A0A367MBX4 | Glutamine--tRNA ligase | 0 | 7.116 | 8 | 4 | 3 | 561 | 63.3 | 83.14 |
| High | L8MK56 | Glutamyl-tRNA(Gln) amidotransferase subunit | 0 | 4.254 | 6 | 1 | 1 | 483 | 51.4 | 58.26 |
| High | A0A2R4BHU9 | Aspartyl/glutamyl-tRNA(Asn/Gln) amidotransferase subunit | 0 | 3.26 | 3 | 3 | 1 | 492 | 54.6 | 50.4 |
| High | A0A485HEP7 | D-aminoacyl-tRNA deacylase | 0 | 3.839 | 4 | 1 | 1 | 468 | 51.9 | 86.76 |
| High | A0A1H0JDB1 | Succinate--CoA ligase | 0 | 25.501 | 17 | 9 | 6 | 388 | 41.5 | 68.46 |
| High | A0A0A8RKE2 | Isoleucine--tRNA ligase | 0 | 42.217 | 14 | 23 | 9 | 943 | 105.4 | 52.02 |
| High | A6V3C2 | Ribonuclease | 0 | 11.6 | 5 | 4 | 3 | 1073 | 119.3 | 49.66 |
| High | A0A086C201 | RNA polymerase | 0 | 8.959 | 6 | 5 | 1 | 620 | 70.1 | 53.84 |
| High | A0A087L9F7 | Alkaline phosphatase family protein | 0 | 7.665 | 12 | 3 | 2 | 269 | 30 | 59.9 |
| High | A0A2R3ISX8 | Bifunctional proteins | 0 | 6.982 | 13 | 3 | 3 | 454 | 48.8 | 83.18 |
| High | A0A1G8LM51 | RNA-binding protein | 0 | 23.925 | 66 | 18 | 6 | 86 | 9.5 | 43.04 |
| High | A0A0A8RIM8 | Chitin-binding protein | 0 | 17.973 | 28 | 12 | 6 | 389 | 41.8 | 46.38 |
| High | A0A0D6INW2 | Ribosome-binding factor | 0 | 10.607 | 32 | 4 | 3 | 130 | 14.7 | 56.04 |
| High | A0A127MQF6 | Ribosome modulation factor | 0 | 5.574 | 21 | 3 | 1 | 72 | 8.4 | 46.14 |
| High | A0A2R3INC8 | Ribosome-recycling factor | 0 | 15.315 | 22 | 10 | 3 | 185 | 20.5 | 86.18 |
| High | A0A1C7BK43 | Periplasmic serine endoprotease | 0.002 | 1.523 | 3 | 1 | 1 | 474 | 50.3 | 65.86 |
| High | A0A3M5DFV1 | ATP-dependent Clp protease ATP-binding subunit | 0 | 13.897 | 8 | 7 | 2 | 452 | 50.1 | 67.32 |
| High | A0A069PZV2 | Enoyl-[acyl-carrier-protein] reductase [NADH] | 0.002 | 1.652 | 6 | 1 | 1 | 265 | 28 | 73.34 |
| High | K7Y4J0 | Alkaline metalloprotease | 0 | 9.962 | 7 | 5 | 2 | 481 | 50.6 | 59.52 |
| High | A0A072ZPJ9 | Metalloprotease | 0 | 7.361 | 6 | 2 | 2 | 449 | 47.8 | 70.46 |
| High | A6VCK8 | ATP-dependent zinc metalloprotease | 0 | 6.179 | 2 | 2 | 1 | 642 | 70.3 | 43.9 |
| High | A0A0A8RQS8 | Zn_protease domain-containing protein | 0 | 10.585 | 20 | 7 | 3 | 221 | 24.3 | 67.76 |
| High | A0A1I1UAD0 | Ferritin-like metal-binding proteins | 0 | 5.342 | 8 | 3 | 1 | 169 | 18.8 | 63.86 |

#### Table S1 (continued)

| Protei | Accession | Description | Exp. q | Sum PEF | Coverag | # PSMs ( | # Unique | # AAs | MW [kDa | Abundanc |
|---|---|---|---|---|---|---|---|---|---|---|
| 1 | 2 | 3 | 4 | 5 | 6 | 7 | 8 | 9 | 10 | 11 |
| Synthases | | | | | | | | | | |
| High | A0A3M5EVT4 | 3-dehydroquinate synthase OS=Pseudomonas aeruginosa O | 0 | 5.761 | 9 | 2 | 2 | 421 | 46.1 | 100 |
| High | A0A069Q8Q1 | 1,4-Dihydroxy-2-naphthoyl-CoA synthase OS=Pseudomonas | 0 | 5.632 | 7 | 1 | 1 | 265 | 28.9 | 100 |
| High | A0A291KBQ0 | S-adenosylmethionine synthase OS=Pseudomonas mendoci | 0 | 5.509 | 7 | 2 | 1 | 396 | 42.7 | 100 |
| High | A0A0A8RA44 | Acetolactate synthase OS=Pseudomonas aeruginosa OX=28 | 0 | 4.099 | 3 | 2 | 1 | 592 | 64.7 | 100 |
| High | A0A0A8RC77 | Anthranilate synthase component 1 OS=Pseudomonas aerug | 0 | 2.933 | 4 | 1 | 1 | 496 | 55.1 | 100 |
| High | A0A3D9EI68 | ATP synthase | 0 | 130.604 | 42 | 77 | 1 | 458 | 49.7 | 37.78 |
| High | A0A0A8RS72 | Hydrogen cyanide synthase | 0 | 33.235 | 29 | 10 | 8 | 464 | 50.4 | 83.46 |
| High | A0A077JN20 | GMP synthase | 0 | 9.213 | 8 | 5 | 3 | 527 | 58.1 | 45.4 |
| High | A0A0A8RJH1 | Glucans biosynthesis glucosyltransferase | 0 | 8.951 | 7 | 4 | 4 | 861 | 97 | 91.06 |
| High | A0A127MNP8 | Argininosuccinate synthase | 0 | 8.785 | 5 | 4 | 2 | 405 | 45.4 | 73.16 |
| High | A0A0F6UIJ7 | Carbamoyl-phosphate synthase large chain | 0 | 8.595 | 3 | 3 | 2 | 1073 | 117.3 | 66.24 |
| High | A0A081HI22 | Tryptophan synthase | 0 | 8.141 | 17 | 2 | 2 | 268 | 28.5 | 84.78 |
| High | A0A0A8RJ65 | PQB biosynthetic 3-oxoacyl-[acyl-carrier-protein] synthase | 0 | 7.385 | 7 | 5 | 2 | 348 | 37.6 | 69.48 |
| High | A0A5K1SNI7 | Chorismate synthase | 0 | 5.676 | 8 | 2 | 2 | 363 | 38.9 | 33.44 |
| High | A0A127MLM1 | Thiazole synthase | 0 | 5.568 | 11 | 3 | 2 | 268 | 28.5 | 42.72 |
| High | A0A6H3G8H9 | Lipopolysaccharide biosynthesis protein | 0 | 5.095 | 7 | 3 | 2 | 662 | 74.5 | 64.84 |
| High | A0A0A8RHR0 | Acetolactate synthase | 0 | 4.525 | 3 | 2 | 1 | 574 | 63 | 90.44 |
| High | A0A0A8RG31 | Dihydrofolate synthase/folylpolyglutamate synthase | 0 | 4.132 | 4 | 2 | 1 | 429 | 46.5 | 61.64 |
| High | A0A3M5DAV0 | Biotin synthase | 0.008 | 1.038 | 3 | 1 | 1 | 390 | 43.3 | 85.78 |
| High | A0A3M5ENA3 | Citrate synthase | 0 | 96.305 | 62 | 53 | 14 | 429 | 47.8 | 51.02 |
| High | A0A0A8RPU8 | Phosphoenolpyruvate synthase | 0 | 55.493 | 22 | 34 | 13 | 791 | 85.8 | 44.82 |
| High | A0A0C6EL94 | Malate synthase | 0 | 45.881 | 29 | 17 | 9 | 725 | 78.6 | 55.46 |
| High | A0A1C7BDZ1 | Pyridoxine 5'-phosphate synthase | 0 | 29.474 | 27 | 18 | 5 | 248 | 27.2 | 44.54 |
| High | A0A086BTL8 | Adenylosuccinate synthetase | 0 | 29.298 | 27 | 16 | 7 | 430 | 46.8 | 45.76 |
| High | A0A069QI07 | Glutathione synthetase | 0 | 21.722 | 25 | 8 | 4 | 317 | 35.7 | 34.52 |
| Oxidoreductase enzymes. | | | | | | | | | | |
| High | A0A5F1BV16 | Re/Si-specific NAD(P)(+) transhydrogenase | 0 | 8.376 | 10 | 3 | 2 | 373 | 38.8 | 100 |
| High | A0A0H2Z7C1 | N-succinylglutamate 5-semialdehyde dehydrogenase | 0 | 5.799 | 4 | 1 | 1 | 488 | 51.5 | 100 |
| High | A0A485EKG2 | Gluconate dehydrogenase | 0 | 5.047 | 3 | 3 | 2 | 1275 | 138.2 | 100 |
| High | A0A7U9F023 | GDP-mannose 6-dehydrogenase | 0.001 | 2.742 | 3 | 1 | 1 | 442 | 48.2 | 100 |

**Note: title of each column is defined as**

**1:** Protein FDR Confidence: Combined.
**2:** Accession.
**3:** Protein description.
**4:** Exp. q-value: Combined.
**5:** Sum PEP Score.
**6:** Coverage [%].
**7:** # PSMs (the total number of identified peptide spectra matched for the protein).
**8:** # of Unique Peptides
**9:** Amino Acid numbers.
**10:** Molecular Weight [kDa]
**11:** Abundances (%).

#### Table S2. Summary of proteomic analysis of D-Exo

| Protei | Accessio | Description | Exp. q- | Sum PEF | Covera | # PSM | # Un | # AAs | MW [kI | Abunda |
|---|---|---|---|---|---|---|---|---|---|---|
| 1 | 2 | 3 | 4 | 5 | 6 | 7 | 8 | 9 | 10 | 11 |
| Fatty acid oxidation and hydrogen peroxide production | | | | | | | | | | |
| High | A0A1C7E | DAO domain-containing protein | 0 | 6.348 | 6 | 2 | 2 | 468 | 52.2 | 44.44 |
| High | A0A2R3Q | D-amino acid dehydrogenase | 0 | 3.474 | 10 | 3 | 1 | 432 | 47 | 100 |
| High | A0A127M | Soluble pyridine nucleotide transhydrogenase | 0 | 112.322 | 57 | 94 | 1 | 464 | 51.3 | 72.72 |
| High | A0A1H0H | 2-hydroxycyclohexanecarboxyl-CoA dehydrogenase | 0.004 | 1.326 | 4 | 2 | 1 | 255 | 26.1 | 67.18 |
| High | A0A080V | CatB-related O-acetyltransferase | 0.002 | 1.726 | 7 | 1 | 1 | 229 | 25.6 | 65.42 |
| High | A0A069Q | Glycerophosphodiester phosphodiesterase | 0 | 23.456 | 26 | 10 | 4 | 240 | 26.9 | 31.98 |
| High | A0A080V | Acetyl-coenzyme A synthetase | 0 | 9.765 | 5 | 3 | 2 | 645 | 71.6 | 35.4 |
| High | A0A1G8E | Glutamate synthase (NADPH/NADH) large chain | 0 | 4.786 | 1 | 1 | 1 | 1482 | 161.9 | 54.66 |
| High | A0A0A8R | Dihydrolipoamide acetyltransferase component of pyruvate ( | 0 | 23.135 | 6 | 10 | 2 | 428 | 45.5 | 42.34 |
| Protein oxidation and activate the Fenton reaction | | | | | | | | | | |
| High | A0A7U0J | AAA family ATPase | 0 | 6.038 | 5 | 3 | 2 | 555 | 57.7 | 80 |
| High | A0A3M5E | Dihydroxy-acid dehydratase | 0.001 | 2.593 | 2 | 2 | 1 | 680 | 72.7 | 39.26 |
| High | A0A3M5D | OMP_b-brl domain-containing protein | 0 | 60.174 | 42 | 27 | 10 | 236 | 25.6 | 53.68 |
| High | A0A3M5D | STN domain-containing protein | 0 | 5.527 | 2 | 2 | 1 | 977 | 105.6 | 71.16 |
| High | A0A1H0I2 | LPS-assembly protein LptD | 0.005 | 1.195 | 1 | 1 | 1 | 912 | 102.8 | 57.06 |
| High | A0A3D9E | Urease accessory protein UreF | 0.007 | 1.136 | 10 | 1 | 1 | 223 | 24.4 | 77.22 |
| High | A0A5F1B | YkgJ family cysteine cluster protein | 0.004 | 1.281 | 15 | 1 | 1 | 223 | 24.8 | 81.88 |
| High | A0A0A8R | UvrABC system protein A | 0 | 13.698 | 7 | 5 | 5 | 1003 | 110.3 | 42.78 |
| High | A0A1G9Y | Sterol carrier protein | 0 | 6.833 | 16 | 1 | 1 | 104 | 11.1 | 100 |
| High | A0A0A8R | Protein PelC | 0 | 7.15 | 12 | 2 | 1 | 172 | 18.6 | 72.7 |
| High | A0A072Z | Amino-acid carrier protein AlsT | 0 | 10.903 | 6 | 2 | 1 | 449 | 47.3 | 31.46 |
| High | W1MFI8 | Lactamase_B domain-containing protein | 0 | 8.994 | 7 | 5 | 2 | 433 | 48.6 | 41.32 |
| High | A0A0A8R | MaoC-like domain-containing protein | 0 | 10.152 | 19 | 4 | 3 | 285 | 31.1 | 39.32 |
| High | A0A080V | Putative periplasmic transport protein | 0 | 8.078 | 11 | 4 | 2 | 319 | 33.6 | 64.1 |
| High | A0A086C | Terminase OS=Pseudomonas aeruginosa VRFPA01 | 0 | 3.309 | 9 | 1 | 1 | 119 | 13.3 | 30.72 |
| unfold DNA, RNA, and proteins | | | | | | | | | | |
| High | A0A0A8R | ATP-dependent RNA helicase RhlB | 0 | 3.444 | 2 | 1 | 1 | 579 | 63.8 | 65 |
| High | A0A072Z | 3-guanidinopropionase | 0 | 5.869 | 6 | 2 | 1 | 318 | 34.2 | 57.04 |
| promote cellular adhesion | | | | | | | | | | |
| High | A0A367L | Aldehyde dehydrogenase family protein (Fragment) | 0.002 | 1.788 | 11 | 2 | 1 | 99 | 11 | 100 |
| High | A0A2R3I | Neisseria PilC beta-propeller domain protein | 0 | 6.646 | 3 | 2 | 1 | 1155 | 126.1 | 100 |
| High | A0A1I1T | Flagellar L-ring protein | 0 | 3.893 | 8 | 1 | 1 | 237 | 24.9 | 100 |
| High | A0A485G | Type 4 fimbrial biogenesis protein PilW | 0 | 11.1 | 11 | 4 | 3 | 625 | 68.7 | 30.4 |
| High | A0A0F6U | Flagellar hook-associated protein 3 | 0 | 121.787 | 60 | 63 | 16 | 439 | 46.8 | 47.68 |
| High | A0A1C7E | Fimbrial assembly protein pilQ | 0 | 33.737 | 21 | 22 | 9 | 714 | 77.4 | 68.1 |
| High | A0A022P | Flagellar basal-body rod protein FlgG | 0 | 31.805 | 39 | 17 | 7 | 261 | 27.7 | 40.98 |
| High | A0A072Z | Fap amyloid fiber secretin | 0 | 5.721 | 8 | 3 | 2 | 421 | 45.7 | 40.02 |
| High | A0A3M5E | Flagellar P-ring protein | 0 | 6.05 | 6 | 3 | 2 | 554 | 57.4 | 90.62 |
| virulence factors | | | | | | | | | | |
| High | A0A7Z0K | Flagellin | 0 | 6.102 | 12 | 5 | 1 | 397 | 40.6 | 100 |
| High | A0A6M5K | B-type flagellin | 0 | 74.411 | 84 | 74 | 7 | 126 | 13.3 | 92.02 |
| iron acquisition | | | | | | | | | | |
| High | A0A0A8R | AMP-binding protein | 0.001 | 2.854 | 2 | 2 | 1 | 555 | 60.1 | 100 |
| High | A0A0H2Z | Heme acquisition protein HasAp | 0 | 6.228 | 11 | 2 | 1 | 205 | 20.9 | 100 |
| High | A6UYT6 | Putative hydroxamate-type ferrisiderophore receptor | 0 | 5.927 | 2 | 1 | 1 | 806 | 88.8 | 100 |
| High | A0A485E | TonB-dependent receptor protein | 0 | 6.662 | 6 | 3 | 3 | 810 | 87.7 | 100 |
| High | A0A7U3T | HAMP domain-containing protein | 0 | 8.311 | 3 | 1 | 1 | 575 | 62.2 | 100 |
| High | A0A3M5E | Protein translocase subunit SecD | 0 | 30.446 | 18 | 11 | 8 | 622 | 67.9 | 32.28 |
| High | A0A0H2Z | Ferripyoverdine receptor | 0 | 268.477 | 54 | 193 | 37 | 815 | 91.1 | 41.62 |
| High | A0A1Y0G | Heme/Hemoglobin uptake outer membrane receptor PhuR | 0 | 165.688 | 46 | 83 | 1 | 764 | 84.6 | 60.88 |
| High | A0A659C | TonB-dependent hemoglobin/transferrin/lactoferrin family rec | 0 | 175.382 | 48 | 90 | 2 | 764 | 84.7 | 42.78 |
| High | A0A0G4Z | Hemin receptor | 0 | 14.348 | 7 | 6 | 4 | 891 | 97.8 | 43.08 |
| High | A0A072Z | Ferric enterobactin receptor | 0 | 8.775 | 7 | 3 | 3 | 746 | 81 | 60.2 |
| High | A0A1C7E | Extracellular heme-binding protein | 0 | 31.419 | 14 | 13 | 9 | 989 | 108.2 | 71.92 |
| High | A0A0A8R | Putative outer membrane protein | 0.001 | 2.824 | 6 | 1 | 1 | 478 | 51.4 | 66.52 |
| High | A0A7U4A | Putative TonB-dependent receptor | 0 | 2.914 | 3 | 1 | 1 | 736 | 81.3 | 68.86 |

#### Table S2 (continuous)

| Protei | Accessio | Description | Exp. q- | Sum PEF | Covera | # PSM | # Un | # AAs | MW [kI | Abunda |
|---|---|---|---|---|---|---|---|---|---|---|
| **1** | **2** | **3** | **4** | **5** | **6** | **7** | **8** | **9** | **10** | **11** |
| Catalyses | | | | | | | | | | |
| High | A0A0A8R | Hydrolase_4 domain-containing protein | 0 | 4.573 | 7 | 2 | 2 | 339 | 38.3 | 100 |
| High | A0A072Z | Nucleotide sugar epimerase/dehydratase WbpM | 0.001 | 2.571 | 2 | 1 | 1 | 665 | 74.3 | 100 |
| High | A0A5E5R | Esterase EstA | 0 | 116.039 | 43 | 74 | 2 | 646 | 69.6 | 74.84 |
| High | A0A7U9F | Transaldolase | 0 | 5.763 | 7 | 3 | 1 | 309 | 33.9 | 34.6 |
| High | A0A7U4A | Soluble pyridine nucleotide transhydrogenase | 0 | 170.703 | 63 | 138 | 5 | 464 | 51.1 | 66.72 |
| High | V6AM38 | Carbamoyl-phosphate synthase small chain | 0.001 | 2.158 | 3 | 1 | 1 | 398 | 43.2 | 100 |
| High | Q9L6C7 | Triacylglycerol acylhydrolase | 0 | 137.528 | 59 | 114 | 3 | 311 | 32.7 | 41.98 |
| High | A0A0H2Z | RND efflux membrane fusion protein | 0 | 40.982 | 31 | 19 | 7 | 370 | 39.1 | 51.02 |
| High | A0A1C7E | Lysin domain-containing protein | 0 | 40.492 | 21 | 22 | 7 | 341 | 37.6 | 75.92 |
| High | A0A077J. | Alpha/beta hydrolase | 0.004 | 1.343 | 4 | 2 | 1 | 275 | 30.4 | 46.26 |
| High | A0A643I( | Protocatechuate 3,4-dioxygenase subunit beta | 0 | 7.402 | 10 | 1 | 1 | 239 | 27.2 | 100 |
| High | A0A0A8R | Uricase | 0 | 3.874 | 6 | 2 | 2 | 494 | 55 | 57.36 |
| High | A0A485E | Signal peptidase I | 0 | 6.5 | 16 | 2 | 2 | 214 | 24 | 40.88 |
| High | A0A1C7E | Lipoprotein | 0 | 6.93 | 7 | 5 | 1 | 210 | 23.7 | 56.8 |
| High | A0A086B | Peptidyl-tRNA hydrolase | 0.004 | 1.248 | 7 | 1 | 1 | 194 | 20.8 | 100 |
| cellular permeability | | | | | | | | | | |
| High | A0A086B | ABC transporter permease | 0.004 | 1.279 | 3 | 2 | 1 | 231 | 24.9 | 54.84 |
| High | A0A3S0I | Autotransporter domain-containing protein | 0 | 104.446 | 21 | 83 | 1 | 991 | 104.4 | 41.58 |
| High | A0A0H2Z | Putative outer membrane receptor protein | 0 | 51.845 | 24 | 19 | 11 | 705 | 79.2 | 33.32 |
| High | Q9HVG7 | POTRA domain-containing proteinX=208964 GN=PA4624 F | 0 | 101.304 | 54 | 57 | 18 | 568 | 63.2 | 37.78 |
| High | A0A7M3/ | Porins | 0 | 71.452 | 59 | 47 | 1 | 204 | 22.9 | 100 |
| High | A0A0A8R | ABC transporter ATP-binding protein/permease | 0.001 | 2.707 | 2 | 1 | 1 | 682 | 74.7 | 100 |
| High | A0A6N4I | Type VI secretion system contractile sheath large subunit | 0 | 23.3 | 18 | 11 | 5 | 491 | 55.5 | 59.14 |
| High | A0A3M5E | ATP-grasp domain-containing protein | 0 | 23.138 | 21 | 9 | 7 | 519 | 59.5 | 64.08 |
| High | A0A080V | Pseudopaline transport outer membrane protein CntO | 0 | 11.172 | 8 | 3 | 3 | 708 | 79 | 55.96 |
| Regulators | | | | | | | | | | |
| High | A0A6A9J | Response regulator | 0 | 3.509 | 9 | 1 | 1 | 212 | 23.2 | 100 |
| High | A0A1H0F | Soluble pyridine nucleotide transhydrogenase | 0 | 77.621 | 33 | 68 | 2 | 464 | 51.1 | 85.7 |
| High | A0A7Y9X | Type I restriction enzyme R subunit | 0.008 | 1.097 | 3 | 1 | 1 | 910 | 103.1 | 100 |
| High | A0A5R1A | TetR family transcriptional regulator | 0.002 | 2.007 | 4 | 1 | 1 | 212 | 24 | 39.9 |
| High | A0A069Q | Protocatechuate 3,4-dioxygenase alpha chain | 0.001 | 2.443 | 7 | 2 | 1 | 201 | 22.8 | 53.08 |
| High | A0A086B | Transcriptional regulator | 0.001 | 2.769 | 5 | 1 | 1 | 225 | 25.5 | 56.6 |
| High | V6AMV0 | Putative methyl-accepting chemotaxis protein | 0 | 13.017 | 3 | 4 | 1 | 859 | 91.1 | 63 |
| High | A0A0A8R | Putative HTH-type transcriptional regulator YdcR | 0 | 14.153 | 11 | 6 | 4 | 500 | 55.3 | 100 |
| High | A0A0D7N | Chemotaxis transducer | 0 | 16.779 | 12 | 4 | 3 | 545 | 58.3 | 71.76 |
| High | A0A0A8R | HIT domain-containing protein | 0 | 4.547 | 7 | 1 | 1 | 209 | 23 | 69.86 |
| High | A0A3M5E | HTH tetR-type domain-containing protein | 0 | 3.801 | 4 | 1 | 1 | 274 | 30 | 100 |
| High | A0A3D9E | Translation initiation factor IF-2 | 0 | 3.756 | 2 | 1 | 1 | 852 | 91.9 | 39.32 |
| High | V6A9P6 | PhoH-like protein | 0 | 2.99 | 5 | 1 | 1 | 363 | 41.2 | 69.26 |

**Note: title of each column is defined as**

**1:** Protein FDR Confidence: Combined.

**2:** Accession.

**3:** Protein description.

**4:** Exp. q-value: Combined.

**5:** Sum PEP Score.

**6:** Coverage [%].

**7:** # PSMs (the total number of identified peptide spectra matched for the protein).

**8:** # of Unique Peptides

**9:** Amino Acid numbers.

**10:** Molecular Weight [kDa]

**11:** Abundances (%).

**References:**

1. Clinic, M. *Drugs and Supplements: Tetracycline (Class) (Oral Route, Parenteral Route)*.
